# Uncoupling the distinct functions of HP1 proteins during heterochromatin establishment and maintenance

**DOI:** 10.1101/2023.04.30.538869

**Authors:** Melissa Seman, Alexander Levashkevich, Ajay Larkin, Fengting Huang, Kaushik Ragunathan

**Affiliations:** Department of Biology, Brandeis University, Waltham, MA 02451 USA

## Abstract

H3K9 methylation (H3K9me) marks transcriptionally silent genomic regions called heterochromatin. HP1 proteins are required to establish and maintain heterochromatin. HP1 proteins bind to H3K9me, recruit factors that promote heterochromatin formation, and oligomerize to form phase-separated condensates. We do not understand how HP1 protein binding to heterochromatin establishes and maintains transcriptional silencing. Here, we demonstrate that the *S.pombe* HP1 homolog, Swi6, can be completely bypassed to establish silencing at ectopic and endogenous loci when an H3K4 methyltransferase, Set1 and an H3K14 acetyltransferase, Mst2 are deleted. Deleting Set1 and Mst2 enhances Clr4 enzymatic activity, leading to higher H3K9me levels and spreading. In contrast, Swi6 and its capacity to oligomerize were indispensable during epigenetic maintenance. Our results demonstrate the role of HP1 proteins in regulating histone modification crosstalk during establishment and identifies a genetically separable function in maintaining epigenetic memory.

## INTRODUCTION

Chromatin organization is critical for maintaining genome stability and regulating gene expression. Nucleosomes are at the core of chromatin structure consisting of DNA wrapped around histone proteins (H2A, H2B, H3, and H4).^1^ Histone modifications mark two classes of chromatin states, transcriptionally active euchromatin and silent heterochromatin. H3K9 methylation (H3K9me) is associated with constitutive heterochromatin regions in organisms ranging from yeast to humans.^2–4^ H3K9me promotes binding of a conserved class of chromatin-associated proteins called heterochromatin protein 1(HP1).^5,6^ HP1 proteins bind to H3K9me via an N-terminal chromodomain (CD) and dimerize through interactions involving a C-terminal chromoshadow domain (CSD).^7,8^ HP1 proteins can recruit effector molecules to heterochromatin, including histone modification enzymes, heterochromatin initiation factors, and nucleosome remodelers.^9,10^ Given the multifaceted properties of HP1 proteins, we do not understand how HP1 proteins are involved in establishing and maintaining epigenetic silencing.^11^

In addition to their recruitment functions, HP1 proteins have structural roles in heterochromatin formation. Based on biochemical studies, HP1 proteins are thought to bind to and bridge adjacent H3K9me nucleosomes.^12^ Additionally, HP1 proteins can oligomerize which facilitates long-range interactions across several kilobases of DNA.^13–15^ *In vivo*, HP1 binding is thought to drive chromatin compaction, thus creating genomic structures that are refractory to RNA polymerase II activity.^16,17^ *In vitro* biophysical measurements support these models wherein HP1 binding to DNA forms protein-DNA complexes that exhibit a stall force of >40pN, representing a barrier to RNA polymerase II elongation.^18^ Although not mutually exclusive, recent studies have also shown that HP1 protein oligomerization forms biological condensates that selectively include heterochromatin proteins while excluding proteins that promote transcription.^19–21^ Because of these wide-ranging biophysical properties of HP1 proteins, all models of heterochromatin are premised on HP1 proteins being essential for transcriptional silencing.^22–25^

In *Schizosaccharomyces pombe (S.pombe),* H3K9me marks constitutive heterochromatin regions, including the pericentromeric repeats flanking centromeres, telomeres, and the mating type locus.^26^ The enzyme Clr4 catalyzes H3K9 methylation, which leads to binding of two conserved HP1 proteins, Swi6 and Chp2.^10,27–29^ Heterochromatin formation in *S.pombe* involves three distinct steps, all of which critically require Swi6 (**Figure 1A**). First, DNA or RNA sequence-dependent *de novo* **initiation** involves site-specific recruitment of Clr4 by RNAi factors. Heterochromatin establishment at the pericentromeric repeats rely primarily on the RNAi pathway. Initial production of small RNAs targeting pericentromeric repeats depends on an abundant class of RNA degradation products called primal RNAs (priRNAs).^30^ Subsequent amplification of the RNAi response depends on the activity of an RNA dependent RNA polymerase, Rdp1 that is recruited to heterochromatin through interactions with Swi6 and an adapter protein, Ers1.^31–33^ In contrast, the mating type locus and telomeres depend both on RNAi and DNA binding proteins (Atf1 and Pcr1 at the mating type locus; Taz1 at the telomeres) to recruit Clr4.^34,35^ Second, **establishment** involves spreading of H3K9me across a chromosomal region of several kilobases.^36,37^ Swi6 interacts with a histone deacetylase, Clr3, and is thought to promote the spreading of H3K9me and transcriptional gene silencing, presumably through its chromatin compaction and phase separation activities.^38^ Third, **maintenance** involves the propagation of H3K9me following DNA replication.^39^ This is an active process that involves the read-write function of the H3K9 methyltransferase, Clr4.^40–42^ Previous studies have shown that in addition to H3K9me, Swi6 can remain bound to heterochromatin at the mating type locus throughout the cell cycle, raising the possibility that an HP1 bound chromatin structure may be essential for epigenetic inheritance.^43^

**Figure 1.**
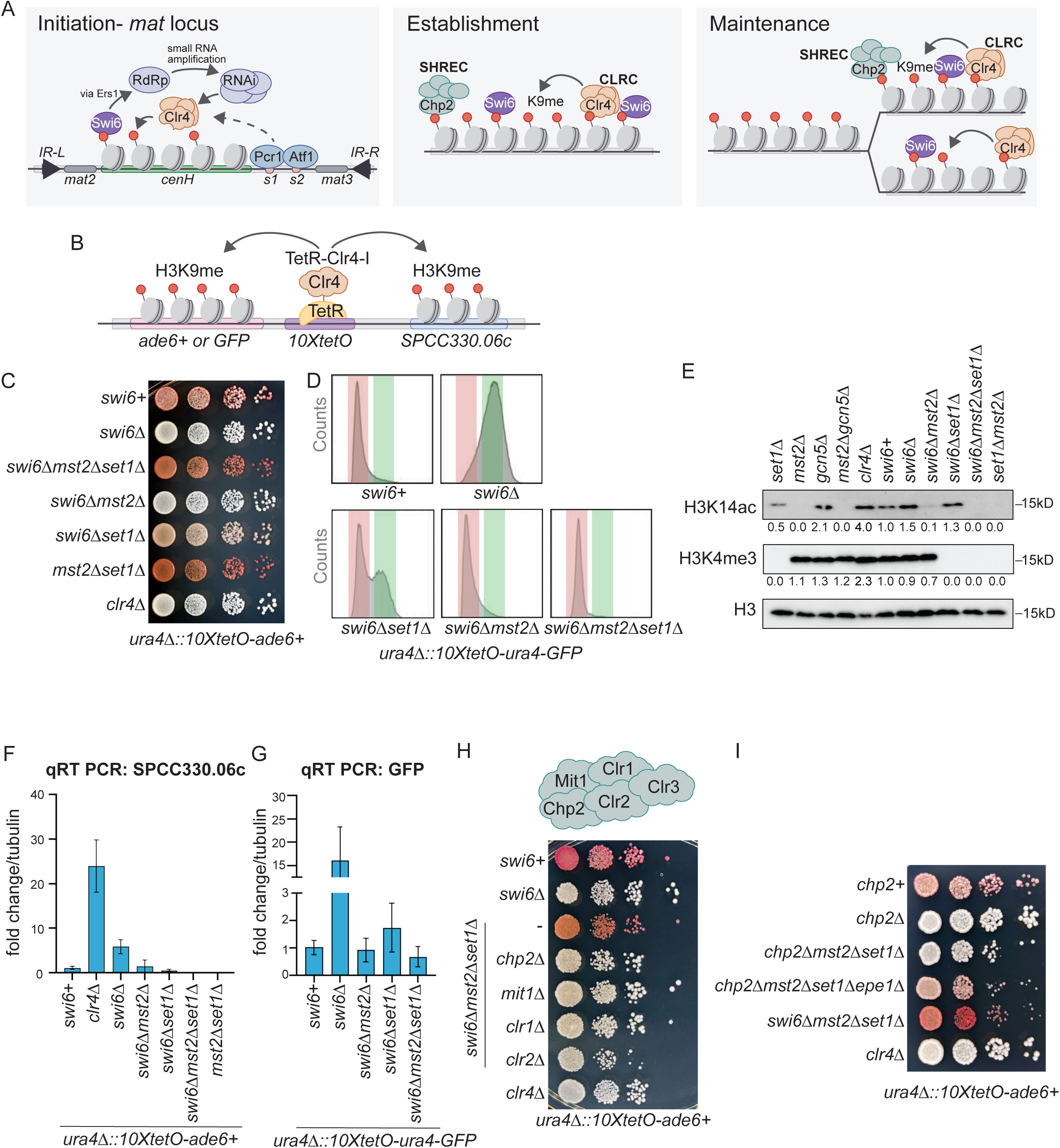
Swi6 can be bypassed in transcriptional gene silencing when either Set1 or Mst2 are deleted. (**A**) Schematic describing the role of Swi6 in the three steps of heterochromatin formation. **Initiation-** DNA or RNA sequence dependent *de novo* initiation involves the site-specific recruitment of Clr4 by RNAi factors. **Establishment-** spreading of H3K9me across a large chromosomal region (several kb) and transcriptional silencing. **Maintenance-** the propagation of H3K9 methylation following DNA replication. (**B**) Schematic of TetR-Clr4-I recruitment to 10XtetO binding sites placed upstream of an *ade6+* or ura4-GFP reporter. (**C**) Silencing assay of *ura4Δ::10XtetO*-*ade6+* reporter in indicated genotypes. Red cells indicate *ade6+* silencing. Cells are plated at 10-fold serial dilutions. (**D**) Flow cytometry to examine GFP expression in *ura4Δ::10XtetO-ura4-GFP* reporter in indicated genotypes. Histograms shifted to the left indicate a GFP-OFF population (red) and those shifted to the right indicate a GFP-ON population (green). (**E**) Western blots comparing levels of H3K14ac and H3K4me3 in indicated genotypes. H3 is shown as a loading control. Quantification is signal in H3K14ac or H3K4me3 blot relative to H3. (**F**) qRT-PCR measurements of RNA levels of *SPCC330.06c* in indicated genotypes. Error bars indicate SD (N=3) (*swi6+* vs *swi6Δ,* p-value =0.0043) (**G**) qRT-PCR measurements of RNA levels of *GFP*. Error bars indicate SD (N=2). (**H**) Silencing assay measuring silencing in the absence of subunits of the SHREC complex. (**I**) Silencing assay measuring silencing in the absence of Chp2. See also Figure S1.

The simple genetic outcome of deleting the major HP1 protein, Swi6 is that it disrupts every step of heterochromatin formation, encompassing initiation, establishment, and maintenance.^44,45^ Hence, it is necessary to leverage genetic tools and cellular contexts that allow us to separately interrogate the role of HP1 proteins in each step of heterochromatin formation. To uncouple the three steps of heterochromatin formation, we utilize an inducible system.^42^ In this system, a site-specific DNA binding, TetR, is fused to the enzymatic SET domain (Clr4 without its chromodomain) of the H3K9 methyltransferase, Clr4 (TetR-Clr4-I). TetR-Clr4-I binding to *10XtetO* sites placed upstream of a reporter gene (*ade6+, ura4+,* or *GFP*) leads to heterochromatin formation and reporter gene silencing. This system captures all of the salient consequences of H3K9me establishment. Additionally, after tetracycline-mediated release of TetR-Clr4-I, we can separately investigate how heterochromatin is maintained after DNA replication. The site-specific recruitment of Clr4 enables the complete bypass RNAi-dependent initiation. Additionally, the TetR-Clr4-I system can be placed at both an ectopic site and the endogenous *mat* locus.^46^ Hence, our inducible system uniquely allows us to test previous models of Swi6 function in both establishing and maintaining epigenetic silencing.

Epigenetic silencing leads to characteristic changes in the repertoire of histone modifications that decorate chromatin. In particular, heterochromatin formation in *S.pombe* leads to a reduction of H3K4 methylation catalyzed by the Set1/COMPASS complex and H3K14 acetylation catalyzed by Mst2 that belongs to the conserved MYST family acetyltransferase complex. In *S.pombe,* Set1 is recruited to promoter regions in a transcription dependent manner where it deposits H3K4me3. Strikingly, its promoter specific H3K4me3 deposition activity during transcription repels heterochromatin invasion and inhibits H3K9me spreading by regulating Clr4 enzymatic activity and disrupting nucleosome stability.^47,48^ Through mechanisms that parallel the protective function of Set1 in the context of euchromatin, Mst2 is recruited to actively transcribed genes and its catalytic activity suppresses ectopic heterochromatin assembly.^49^ Deleting Mst2 also has a remarkable influence on heterochromatin stability.^50^

Surprisingly, we discovered that Swi6 could be completely bypassed at both an ectopic site and the endogenous *mat* locus during the establishment of transcriptional gene silencing in *S. pombe*. Bypassing Swi6 required deleting the H3K4 methyltransferase, Set1, and the H3K14 acetyltransferase, Mst2. In contrast, Swi6 was essential for maintenance. Mutations that disrupt Swi6 oligomerization resulted in a loss of epigenetic inheritance. Our synthetic approach enabled us to define how Swi6 shifts the equilibrium of histone post-translational modifications towards a heterochromatin-like state that is necessary to maximize Clr4 enzymatic activity. Our studies reveal how HP1 proteins regulate histone modification crosstalk and have an essential structural role in epigenetic inheritance.

## RESULTS

### Swi6 can be bypassed in transcriptional gene silencing when either Set1 or Mst2 are deleted

To evaluate the effects of Swi6 downstream of H3K9 methylation, we used a synthetic approach where we tether a TetR-Clr4-I fusion protein at *10XtetO* binding sites placed upstream of a reporter gene (**Figure 1B**). As previously shown, TetR-Clr4-I recruitment induced heterochromatin and silenced an *ade6+* reporter placed downstream of the *10XtetO* binding sequences **(Figure 1C**, *swi6+*). When the *ade6+* reporter was silenced, cells turned red when plated on media containing low adenine.^51^ In contrast, when *ade6+* was expressed, cells remained white. As expected, deleting Swi6 in cells where we tether Clr4 led to the loss of epigenetic silencing (**Figure 1C**, *swi6Δ*).

Given the unique characteristics of histone modifications associated with euchromatin versus heterochromatin, we wondered if altering the balance of histone modifications associated with active transcription could restore silencing in cells lacking Swi6. To test whether we could reverse transcriptional activation and restore epigenetic silencing, we deleted Mst2, the catalytic subunit of a MYST acetyltransferase complex associated with H3K14 acetylation, or Set1, the catalytic subunit of the Set1C H3K4 methyltransferase complex, in *swi6Δ* cells (**Figure 1C**, *swi6Δmst2Δ* or *swi6Δset1Δ)*. We observed either no rescue upon deleting Mst2 (*swi6Δmst2Δ*) or a limited rescue of *ade6+* silencing upon deleting Set1 (*swi6Δset1Δ)*. In contrast, deleting both Set1 and Mst2 had an additive effect causing colonies to turn red consistent with the full restoration of *ade6+* silencing in *swi6Δ* cells (**Figure 1C**, *swi6Δset1Δmst2Δ*).

To determine whether our results were generalizable to other promoters and reporter systems, we investigated silencing in strains with a *ura4-GFP*.^42^ Silencing of the GFP reporter correlates with a low fluorescence state (GFP-OFF), and GFP expression correlates with a high fluorescence state (GFP-ON). As we have previously shown, TetR-Clr4-I recruitment to *10XtetO* binding sites placed upstream of a *ura4-GFP* reporter led to low GFP expression (**Figure 1D**, *swi6+)*. Deleting Swi6 caused cells in a GFP-OFF state to shift to a GFP-ON state (**Figure 1D**, *swi6Δ*). In the GFP reporter strains, deleting Set1 in the absence of Swi6 (**Figure 1D**, *swi6Δset1Δ*) partially rescued silencing while deleting Mst2 resulted in a full rescue of GFP silencing (**Figure 1D**, *swi6Δmst2Δ*). Combining both deletions in the absence of Swi6 (**Figure 1D**, *swi6Δset1Δmst2Δ*) restored silencing similarly to what we previously observed in strains with an *ade6+* reporter. Although the two reporters behave differently, likely due to differences in their promoter sequences, deleting both Set1 and Mst2 consistently produced an additive effect. Furthermore, we performed Western blot to analyze the H3K14ac and H3K4me3 in each of these genotypes (Figure 1E). As expected H3K14ac was substantially diminished in *mst2Δ* cells and H3K4me3 was completely absent in *set1Δ* cells. Deleting *gcn5Δ* in *mst2Δ* cells produced an additive effect and completely abolished H3K14ac consistent with previous studies.^52^ We also observed unexpected cross-talk between the absence of H3K4me3 and H3K14ac since cells lacking *set1Δ* exhibited lower genome-wide H3K14ac levels potentially explaining the partial rescue of silencing we observed in *swi6Δset1Δ* strains.

We corroborated the results of our silencing assays by measuring transcript levels using reverse transcription followed by quantitative PCR (qRT-PCR). In the *ade6+* reporter strains, we performed qRT-PCR analysis on a gene proximal to the TetR-Clr4-I binding site, *SPCC330.06c* (strains containing the *ade6+* reporter have an endogenous *ade6-M210* allele which confounded direct qRT-PCR analysis).^53^ Consistent with the results of our silencing assay, *swi6Δset1Δmst2Δ* cells fully rescued the silencing defect of *SPCC330.06c* observed in *swi6*Δ cells (**Figure 1F**). RNA levels in *swi6Δmst2Δset1Δ* are further reduced approximately 10-fold lower in expression relative to *swi6+.* We observed an intermediate level of *SPCC330.06c* expression in *swi6Δmst2Δ* (similar gene expression in *swi6+)* and *swi6Δset1Δ* (2-fold reduction in silencing relative to *swi6+*). qRT-PCR of the GFP reporter also matched the trends we observed based on fluorescence intensity measurements. GFP RNA levels increased in *swi6*Δ cells, but silencing was completely rescued in *swi6Δmst2Δ* and *swi6Δmst2Δset1Δ* relative to *swi6+* cells (**Figure 1G**). Intermediate levels of silencing were observed in *swi6Δset1Δ* (2-fold higher than *swi6+*).

We also confirmed that deleting other subunits of the Set1 complex subunits (Ash2, Swd1, Swd2, Swd3, and Sdc1) instead of Set1 (*swi6Δmst2Δ+*Set1 complex deletion) or other subunits of the Mst2 complex (Nto1, Eaf6, Ptf2, and Pdp3) instead of Mst2 (*swi6Δset1*Δ+Mst2 complex deletion) produced a similar rescue of silencing.^50,54^ All deletions except for *swi6Δmst2Δswd3Δ* rescued silencing in the *ade6+* reporter colorimetric assay (**Figure S1A-B**). We confirmed this result using qRT-PCR assays of the endogenous *SPCC330.06c* gene (**Figure S1C-D**). Finally, we confirmed that deleting proteins involved in the RNAi pathway, an endoribonuclease, Dcr1 (*swi6Δmst2Δset1Δdcr1Δ)*, or the small RNA binding protein, Ago1 (*swi6Δmst2Δset1Δago1Δ)*, did not affect silencing.^44^ Cells remained red after the deletion of Dcr1 and Ago1, suggesting that the bypass of Swi6 at the ectopic locus upon deleting Set1 and Mst2 is distinct from any role that Swi6 may have in RNAi dependent initiation (**Figure S1E**).

We measured two characteristic features of heterochromatin in *swi6Δmst2Δset1Δ* cells. First, we measured RNA polymerase II occupancy with ChIP-qPCR of *SPCC330.06c* using an antibody that targets the large subunit of the RNA polymerase II complex (Rpb1). Rpb1 occupancy increased upon deleting Swi6 consistent with the loss of transcriptional gene silencing.^29^ Rpb1 occupancy was restored to near Swi6 wild-type levels in *swi6Δmst2Δset1Δ* cells (**Figure S1F**). Second, we performed histone turnover assays using recombination-induced tag exchange (RITE) which measures the incorporation of new histone H3 tagged across the genome.^55^ We measured low histone turnover in *swi6+* cells. Histone turnover increased in *swi6Δmst2Δ* cells suggesting that despite its well-established role^56^, deleting Mst2 alone was not sufficient in suppressing histone turnover. However, *swi6Δmst2Δset1Δ* cells exhibit substantially reduced histone turnover matching what we observed in the case of successful heterochromatin establishment in *swi6+* cells (**Figure S1G**).

Following H3K9 methylation deposition, epigenetic silencing additionally depends on distinct activities of the SHREC complex, which consists of the second HP1 protein, Chp2, a chromatin remodeler, Mit1, and a histone deacetylase, Clr3 (Figure S1H).^28,29,57–59^ We deleted subunits of the SHREC complex (Mit1, Clr1, Clr2, Chp2) in *swi6Δmst2Δset1Δ* cells and found that silencing was disrupted in each case (**Figure 1H**). These observations suggest that even when Swi6 is bypassed during heterochromatin establishment, this process still requires the well-established roles of downstream chromatin factors such as the SHREC complex. We were unable to delete Clr3 in *swi6Δmst2Δset1Δ*, likely because of a negative genetic interaction resulting in a synthetic lethal phenotype.

Finally, we asked if Chp2, a second HP1 protein and component of SHREC, can also be bypassed by deleting Set1 and Mst2. Consistent with previous work that demonstrates a separate and nonoverlapping function for Chp2 and Swi6, deleting Mst2 or Set1 or both proteins does not restore epigenetic silencing in *chp2Δ* cells (**Figure 1I**). Epe1, the putative H3K9 demethylase, is recruited to chromatin through an interaction with Swi6. In *chp2Δmst2Δset1Δ* cells we wondered if the recruitment of Epe1 by Swi6 was preventing heterochromatin establishment. Indeed, upon deleting Epe1 we found that Chp2 could also be successfully bypassed in establishing heterochromatin (*chp2Δmst2Δset1Δepe1Δ,* **Figure 1I**).

### Swi6 independent silencing leads to increased levels of H3K9me2 and H3K9me3

We performed differential gene expression analysis relative to fully derepressed *clr4*Δ cells, focused on genes within a ∼35kb region surrounding the TetR-Clr4-I binding site. In *swi6+* cells, several genes within the 35kb window exhibit a 2-4 fold decrease in gene expression relative to *clr4*Δ cells (**Figure 2A**). Deleting Swi6 (*swi6Δ)* disrupted silencing, but silencing was fully restored in *swi6Δmst2Δset1Δ* cells across the same 35kb region. In addition, we observed a partial rescue of silencing of many genes proximal to the TetR-Clr4-I tethering site upon deleting only Set1 or Mst2 (*swi6Δset1Δ* or *swi6Δmst2Δ*). Hence, silencing can spread across a large chromosomal region even in the absence of Swi6 following TetR-Clr4-I mediated H3K9me nucleation.

**Figure 2.**
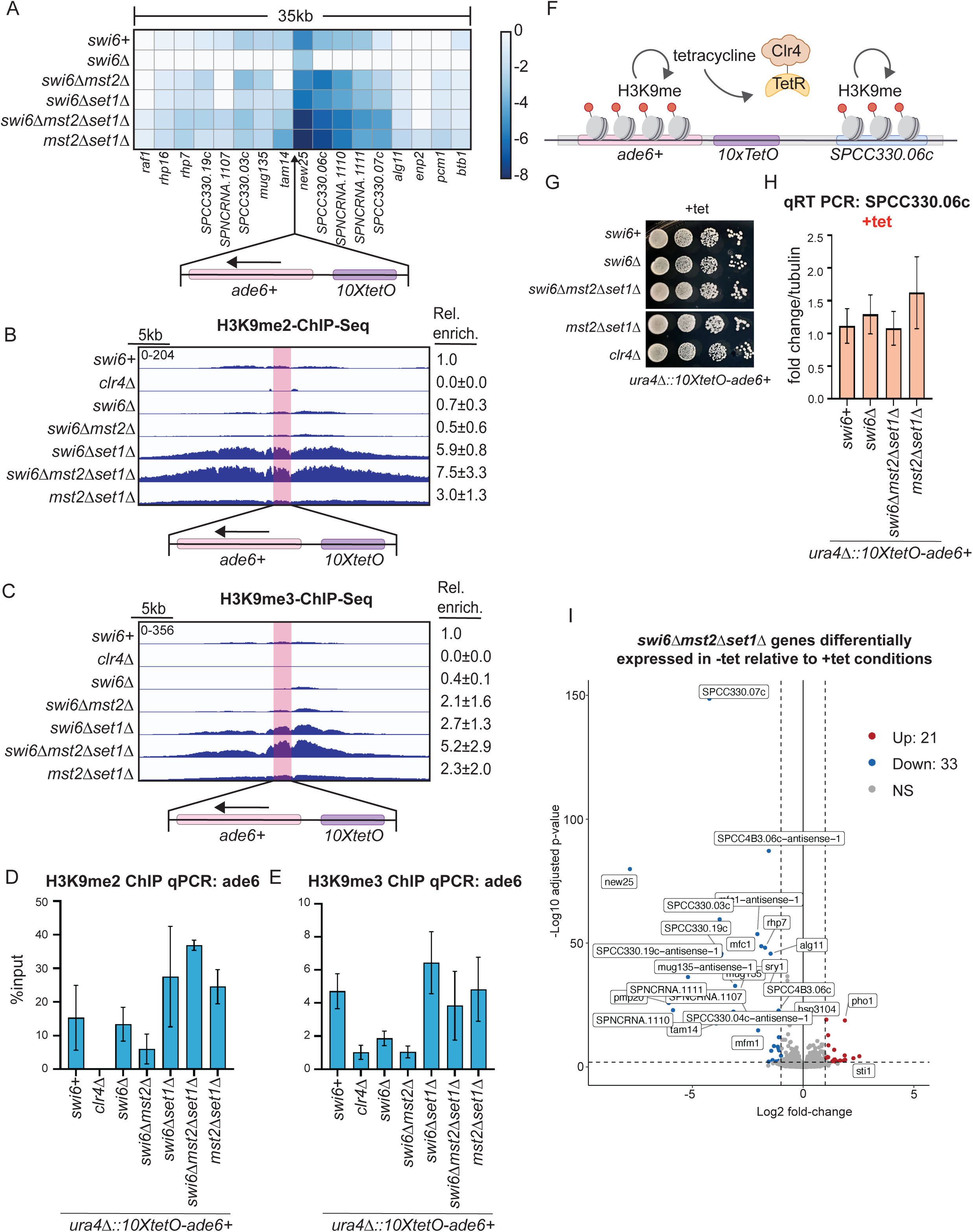
Silencing relies on H3K9me but not Swi6 when Set1 and Mst2 are deleted. (**A**) RNA-seq data examining expression levels the indicated genotypes in a 35kb window surrounding the TetR-Clr4-I binding site. Changes in expression are log2 fold relative to *clr4Δ* cells (N=3-4). (**B,C**) ChIP-seq of H3K9me2 (B) and H3K9me3 (C) surrounding the TetR-Clr4-I binding site in the indicated genotypes. The *10xtetO-ade6+* at the *ura4* locus is highlighted in red. All samples are normalized to input and reads per kilobase per million (RPKM). Relative enrichment is number of reads relative to *swi6+* with SD (N=2). (**D, E**) ChIP-qPCR to measure H3K9me2 (D) or H3K9me3 (E) at the *ade6+* reporter. Error bars indicate SD (N=3). (**F**) Schematic of TetR-Clr4-I release with tetracycline addition. (**G**) Silencing assay of *ura4Δ::10XtetO*-*ade6+* reporter in indicated genotypes on +tet media.(**H**) qRT-PCR measuring RNA levels at *SPCC330.06c* in the indicated genotypes in +tet. Error bars indicate SD (N=3). (**I**) Volcano plots comparing genes downregulated in -tet relative to +tet conditions in *swi6Δmst2Δset1Δ* cells. Genes proximal to the 10xtetO-ade6+ reporter are labelled in the volcano plot. See also Figure S2.

We used chromatin immunoprecipitation followed by sequencing (ChIP-seq) to determine the extent of H3K9me2 and H3K9me3 spreading across the locus where TetR-Clr4 is tethered. Deleting Swi6 led to a substantial loss of H3K9me2 and H3K9me3 across the ectopic locus despite TetR-Clr4-I being bound (*swi6Δ*, **Figure 2B-C**). Hence, the ectopic locus is highly sensitive to the presence of Swi6 for establishing and spreading H3K9 methylation. When Mst2 is deleted, there was no increase in H3K9me2 and only a slight increase in H3K9me3 across the entire region (**Figure 2B-C**, *swi6Δmst2Δ*). However, when Set1 only or Set1 and Mst2 were simultaneously deleted, both H3K9me2 and H3K9me3 were increased 3-8-fold higher. We validated our ChIP-seq results using ChIP-qPCR and observed a 3-fold increase in H3K9me2 and similar levels of H3K9me3 in *swi6Δmst2Δset1Δ* relative to *swi6+* with the *ade6+* reporter (**Figure 2D-E**).

### Swi6 independent silencing requires sequence-specific Clr4 tethering and H3K9 methylation

One possibility is that depleting the epigenome of active histone modifications disrupts transcription on a genome-wide scale leading to H3K9 methylation-independent silencing. To determine if the mechanism of Swi6 independent silencing in *swi6Δmst2Δset1Δ* strains still required H3K9 methylation, we added tetracycline to release TetR-Clr4-I from its binding site^42^ and measured subsequent changes in RNA levels (**Figure 2F**). In the absence of Clr4 tethering, we expected the complete loss of H3K9 methylation and a concomitant loss of silencing. Consistent with our hypothesis, adding tetracycline disrupted the silencing of the *ade6+* reporter across all genotypes, causing cells that were initially red to turn white on +tet containing medium (**Figure 2G**). Additionally, qRT-PCR confirmed that RNA transcript levels of *SPCC330.06c* returned to levels comparable to that of cells with functional Mst2 and Set1 and no Clr4 tethering (**Figure 2H**).

We compared RNA-Seq data of cells grown in +tet to cells grown without tet. We observed that 33 genes were specifically downregulated in *swi6Δmst2Δset1Δ* when TetR-Clr4-I was bound (-tet). Of these 33 genes, 24 were within a 200kb window surrounding the TetR-Clr4-I binding site (**Figure 2I**). In addition, we observed a similar number of genes silenced in a TetR-Clr4-I dependent manner in *swi6+mst2Δset1Δ* (**Figure S2A**). Taken together, these results suggest that Swi6 independent silencing is dependent on TetR-Clr4-I recruitment leading to H3K9 methylation rather than spurious silencing effects due to the deletion of Set1 and Mst2.

### The requirement of Swi6 for transcriptional gene silencing at the mating type locus can be bypassed upon deleting Set1 and Mst2

Swi6 plays an essential role in RNAi-mediated heterochromatin initiation, which serves as the primary mechanism for H3K9 methylation deposition at the pericentromeric repeats and the *mat* locus. RNAi-dependent initiation at the *mat* locus can be bypassed in strains where the RNAi-initiating sequence, *cenH*, is replaced with nine copies of the *tetO* binding sites.^46^ This system recapitulates key heterochromatin features at the *mat* locus, including *cis* inheritance of ON and OFF epigenetic states. Hence, TetR-Clr4-I tethering at the *mat* locus allows us to expand our investigation of the requirement for Swi6 in transcriptional gene silencing to endogenous loci in *S. pombe*.

We used phenotypic assays based on the silencing of a *ura4+* reporter placed downstream of the TetR-Clr4-I binding site. Cells can grow on media containing 5-fluoroorotic acid (FOA) only when *ura4+* is silenced. As previously shown, we observed silencing of the *ura4+* reporter in *swi6+* cells (**Figure 3A**, *swi6+*). Deleting Swi6 caused a loss of reporter gene silencing, leading to cells that failed to grow on FOA-containing media (**Figure 3A**, *swi6Δ*). Deleting Mst2, Set1, or both proteins led to a rescue of silencing of *ura4+* reporter compared to *swi6Δ* cells as evidenced by a rescue of growth on FOA-containing media (**Figure 3A**). We quantified the results of our growth assays by measuring *ura4+* levels using qRT-PCR measurements. Indeed, deleting Set1 or Mst2 reduces *ura4+* gene expression by half. Deleting both Set1 and Mst2 had an additive effect that reduced gene expression 5-fold relative to *swi6Δ* and achieved levels comparable to wild-type *swi6+* cells (**Figure 3B**). We used ChIP-seq to map H3K9me2 and H3K9me3 patterns across the *mat* locus. We found similar levels of H3K9me2 and H3K9me3 in wild-type and *swi6Δ* cells, likely because of the RNAi-independent recruitment of Clr4 through DNA binding proteins, Pcr1 and Atf1 (**Figure 3C**).^34^ In contrast, H3K9me2 and H3K9me3 spreading increased substantially across the *mat* locus relative to wild-type cells with either *swi6Δmst2Δ* or *swi6Δset1Δ*. Notably, deleting both Set1 and Mst2 also produced an additive effect on both H3K9me2 and H3K9me3 spreading across the *mat* locus.

**Figure 3.**
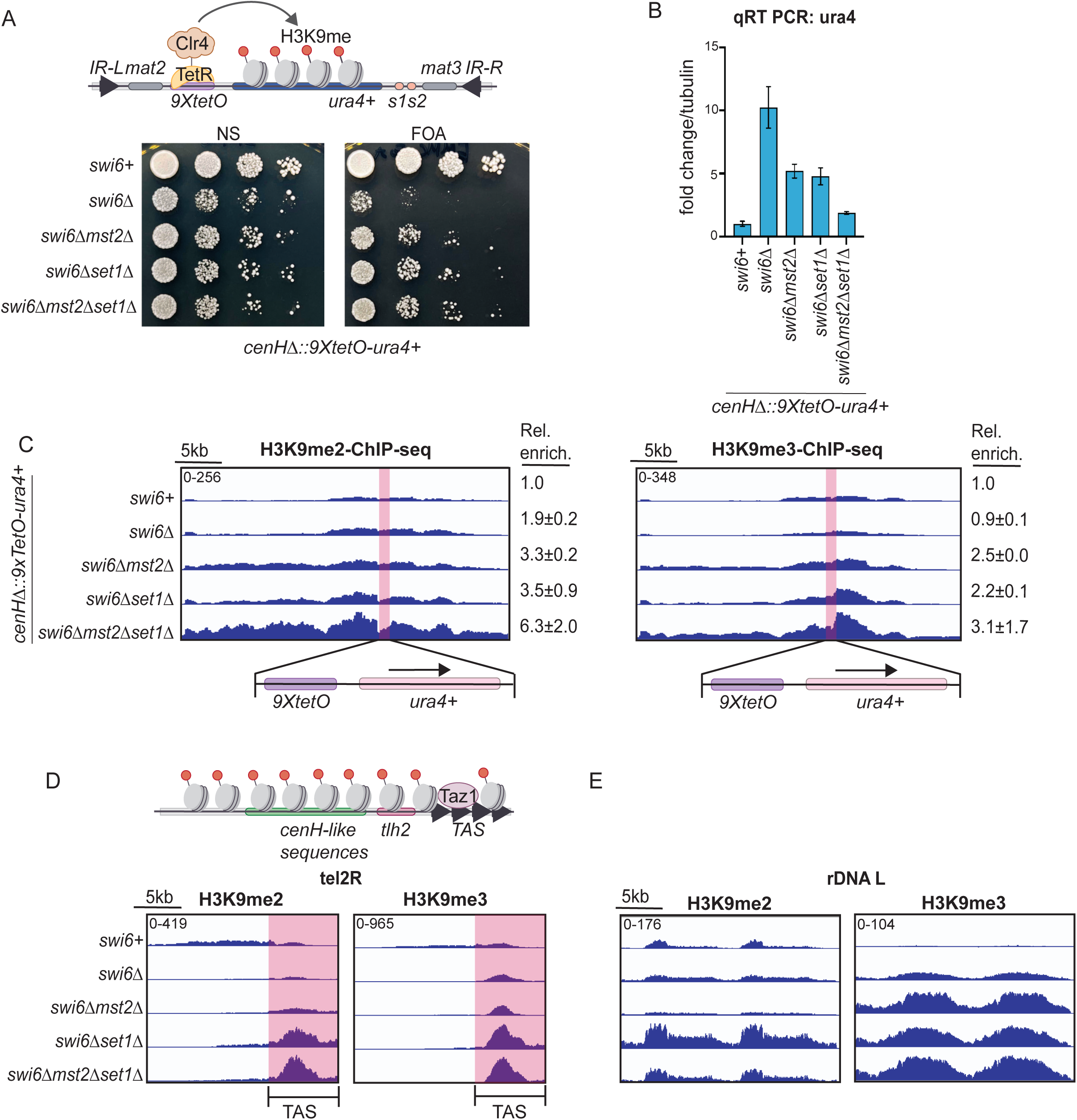
Swi6 is bypassed in heterochromatin formation when two writers of active transcription, Set1 and Mst2, are deleted at an endogenous locus. (**A**) Inducible heterochromatin establishment at the mat locus. 9XtetO-ura4+ gene replaces the RNAi dependent cenH initiating sequence at the mating type locus. Silencing of ura4+ reporter viaTetR-Clr4-I binding in the indicated genotypes. Cells grow on FOA only when ura4+ is silenced. (**B**) qRT-PCR to measure RNA levels of the ura4+ reporter in the indicated genotypes. Error bars indicate SD (N=2). (**C**) ChIP-seq of H3K9me2 and H3K9me3 at the mating type locus. The 9xtetO-ura4+ locus is highlighted in red. All samples are normalized to input and reads per kilobase per million (RPKM). Relative enrichment is number of reads relative to swi6+ with SD (N=2). (**D**) Schematic of the telomere consisting of TAS sequences and RNAi dependent cenH like sequences. ChIP-seq of H3K9me2 and H3K9me3 at the telomere. The telomere associated sequence (TAS) is highlighted in red.(**E**) ChIP-seq of H3K9me2 and H3K9m3 at rDNAL. See also Figure S3.

### TetR-Clr4 tethering is required to bypass the requirement for Swi6 at RNAi-dependent heterochromatin sites

To test whether we could bypass the requirement for Swi6 if RNAi-dependent initiation is intact *(cenH* at the mating type locus and dg/dh repeats at the pericentromeric repeats), we deleted Swi6 in strains that do not have TetR-Clr4-I mediated initiation. Consistent with the essential role of Swi6 in RNAi-mediated heterochromatin formation, we were unable to rescue silencing of a *ura4+* reporter at the *mat* locus in strains that retain the centromere homology region, *cenH* or silencing of an *ade6+* reporter at the pericentromeric repeats (**Figure S3A-B**).^60,61^ We observed a loss of silencing across both reporter systems upon deleting Swi6 and no rescue of silencing upon deleting Set1 and Mst2 in conjunction with Swi6.

At the telomere ends, initiation of heterochromatin formation relies on the DNA binding protein Taz1 rather than RNAi.^35^ The mechanism of H3K9 methylation deposition at the rDNA locus is poorly understood, but it is known to occur through an RNAi-independent pathway.^62^ Our ChIP-seq data showed H3K9me2 and H3K9me3 levels were increased in *swi6Δmst2Δset1Δ* cells relative to *swi6+* or *swi6Δ* cells at TAS sequences but not *cenH*-like sequences (**Figure 3D**). Furthermore, at the rDNA locus, H3K9me2 and H3K9me3 levels were also increased in *swi6Δmst2Δset1Δ* relative to *swi6+* or *swi6Δ* cells (**Figure 3E**). Therefore, we concluded that the requirement for Swi6 can be bypassed at other endogenous sites of heterochromatin formation by deleting Set1 and Mst2 if heterochromatin establishment is RNAi independent or if we can subvert the RNAi dependence at specific loci via TetR-Clr4 tethering.

### Deleting Set1 and Mst2 promotes Clr4 hyperactivity

Previous studies have shown that H3K4me3 inhibits Clr4 enzymatic activity while H3K14 deacetylation, followed by ubiquitination, promotes Clr4 enzymatic activity.^48,63^ This suggests the potential for crosstalk between H3K4me3 and H3K14ub in regulating Clr4 enzymatic activity. Therefore, we wondered if deleting Set1 and Mst2 primes chromatin by creating an ideal chromatin substrate for Clr4 *in vivo,* thus enabling us to bypass the requirement for Swi6.

Consistent with characteristic signatures of Clr4 hyperactivity, we observed an increase of H3K9me at rDNA and telomeres in all *swi6Δ* rescue strains (**Figure 3D-E**, *swi6Δmst2Δ, swi6Δset1Δ, swi6Δmst2Δset1Δ*).^64^ We also observed H3K9me at meiotic genes such as *mei4* and facultative H3K9me islands such as *ssm4* in *swi6Δset1Δ* and *swi6Δmst2Δset1Δ* cells (**Figure 4A**). In addition, we also observed a *de novo* peak of H3K9me at *clr4* which has been observed under conditions of heterochromatin misregulation (**Figure 4B**).^56^

**Figure 4.**
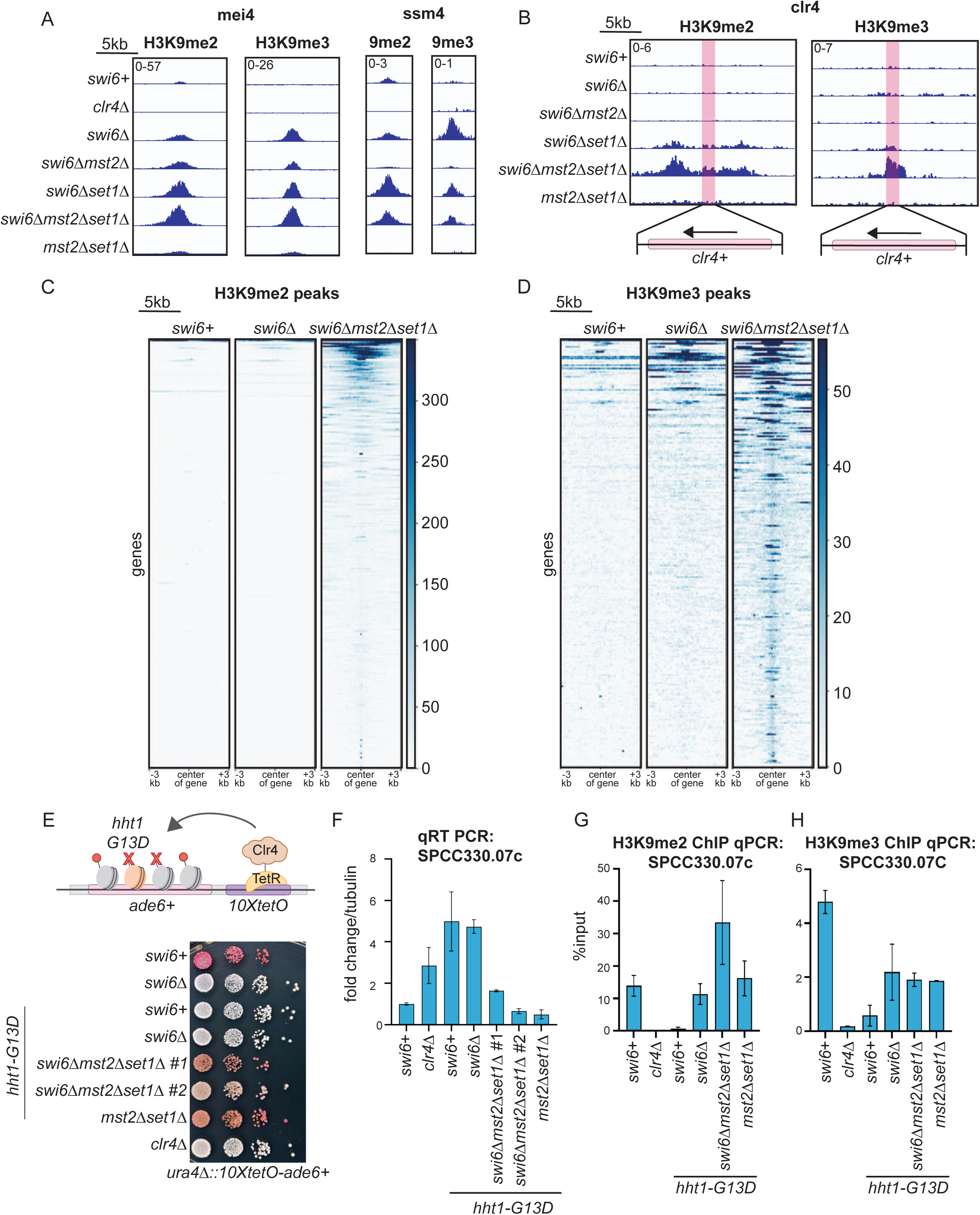
Deleting Set1 and Mst2 promotes Clr4 hyperactivity. (**A**) ChIP-seq of H3K9me2 and H3K9me3 at a meiotic gene *mei4* and a facultative H3K9me island *ssm4*. All samples are normalized to input. (**B**) ChIP-seq of H3K9me2 and H3K9me3 at *clr4*. All samples are normalized to input. (**C, D**) Genome wide heat map of H3K9me2 peaks (C) or H3K9me3 peaks (D). Peaks were first called in *swi6Δmst2Δset1Δ* and compared to H3K9me peaks at the same genomic loci in *swi6+* and *swi6Δ*. (**E**) Schematic showing the genetic outcome of replacing a single histone H3 allele with H3-G13D (*hht1-G13D*). Silencing assay of *ura4Δ::10XtetO*-*ade6+* reporter in the indicated genotypes. (**F**) qRT-PCR to measure gene expression at *SPCC33.07c* in the indicated genotypes. Error bars indicate SD (N=2). (**G, H**) ChIP-qPCR to measure H3K9me2 (G) or H3K9me3 (H) at *SPCC330.07c*. Error bars indicate SD (N=2).

We performed genome-wide analysis on all H3K9me2 and H3K9me3 peaks across *swi6+, swi6Δ,* and *swi6Δset1Δmst2Δ* cells. We called peaks found in *swi6Δmst2Δset1Δ* and compared their genomic coordinates to both *swi6+* and *swi6Δ.* We saw an increase of ectopic H3K9me2 and H3K9me3 peaks at several locations in the genome in *swi6Δmst2Δset1Δ* that were not found in *swi6+* or *swi6Δ* (**Figure 4C-D**). These results support the hypothesis that Clr4 is hyperactive when Set1 and Mst2 are deleted in the absence of Swi6.

To validate our model of enhanced Clr4 activity upon deleting Set1 and Mst2, we engineered strains with a histone H3-G13D (*hht1-G13D*) mutation by editing one of three histone H3 copies in *S.pombe*. *hht1-G13D* is a dominant negative allele that impairs Clr4 enzymatic activity, acts in a dose-sensitive manner, and disrupts heterochromatin establishment by altering H3K9 methylation density.^65^ In strains with the ectopic *10XtetO-ade6+* reporter cells that were initially red turned white upon expressing one allele of *hht1-G13D* (**Figure 4E**). However, deleting Set1 and Mst2 in a *swi6+* or *swi6Δ* background restored silencing in *hht1-G13D* expressing cells, as indicated by the appearance of red colonies across multiple independent biological replicates. We quantified RNA levels using qRT-PCR and found that *SPCC330.06c* was not sensitive to the *hht1-G13D* mutation across different strain backgrounds (data not shown). Instead, we measured expression levels of a second proximal gene, *SPCC330.07c,* in which case silencing was disrupted in cells expressing one allele of *hht1-G13D* but fully restored in *swi6Δmst2Δset1Δhht1-G13D* cells where we observed an approximately 4-fold reduction in mRNA levels (**Figure 4F**). We additionally performed ChIP-qPCR experiments to measure H3K9me2 and H3K9me3 levels in *hht1-G13D* expressing cells. ChIP-qPCR revealed that both H3K9me2 and H3K9me3 were significantly reduced by the *hht1-G13D* mutation in *swi6+* cells (*swi6+, hht1-G13D*, **Figure 4G-H**). However, deleting both Set1 and Mst2, restored high H3K9me2 and H3K9me3 relative to wild-type cells (*swi6Δmst2Δset1Δ, hht1-G13D*). Hence, although H3-G13D attenuates Clr4 enzymatic activity and disrupts silencing, deleting Set1 and Mst2 rescues this defect.

### Swi6 is essential for epigenetic inheritance of H3K9 methylation

Given that we could successfully bypass Swi6 during heterochromatin establishment, we wanted to test whether the same genetic conditions could allow for the bypass of Swi6 in the maintenance of epigenetic silencing. Adding tetracycline triggers the dissociation of the TetR-Clr4-I initiator, which enables us to measure H3K9me maintenance separately from its sequence-dependent establishment. In strains where we established heterochromatin at an ectopic locus, epigenetic inheritance only occurs in the absence of the putative H3K9 demethylase, Epe1 (**Figure 5A**, *epe1Δ*).^40,42,66^ Therefore, we tested whether H3K9me and epigenetic silencing could persist in *swi6Δmst2Δset1Δepe1*Δ strains. As expected, deleting Epe1 in *swi6Δmst2Δset1Δ* did not affect the establishment of *ade6+* silencing, as indicated by red colonies on media lacking tetracycline (**Figure 5A**, *epe1Δswi6Δmst2Δset1Δ)*. However, unlike *swi6+epe1Δ* cells that successfully maintain epigenetic silencing and exhibit a sectored phenotype when plated on +tetracycline containing medium, *swi6Δmst2Δset1Δepe1Δ* cells turned white on tetracycline (**Figure 5A**). In addition, *swi6Δmst2Δset1Δepe1Δ* cells exhibited a complete loss of H3K9me2 and H3K9me3 at the *ade6+* reporter (**Figure 5B-C**). In contrast, cells that have *swi6+* retained substantial levels of H3K9me2 and H3K9me3.

**Figure 5.**
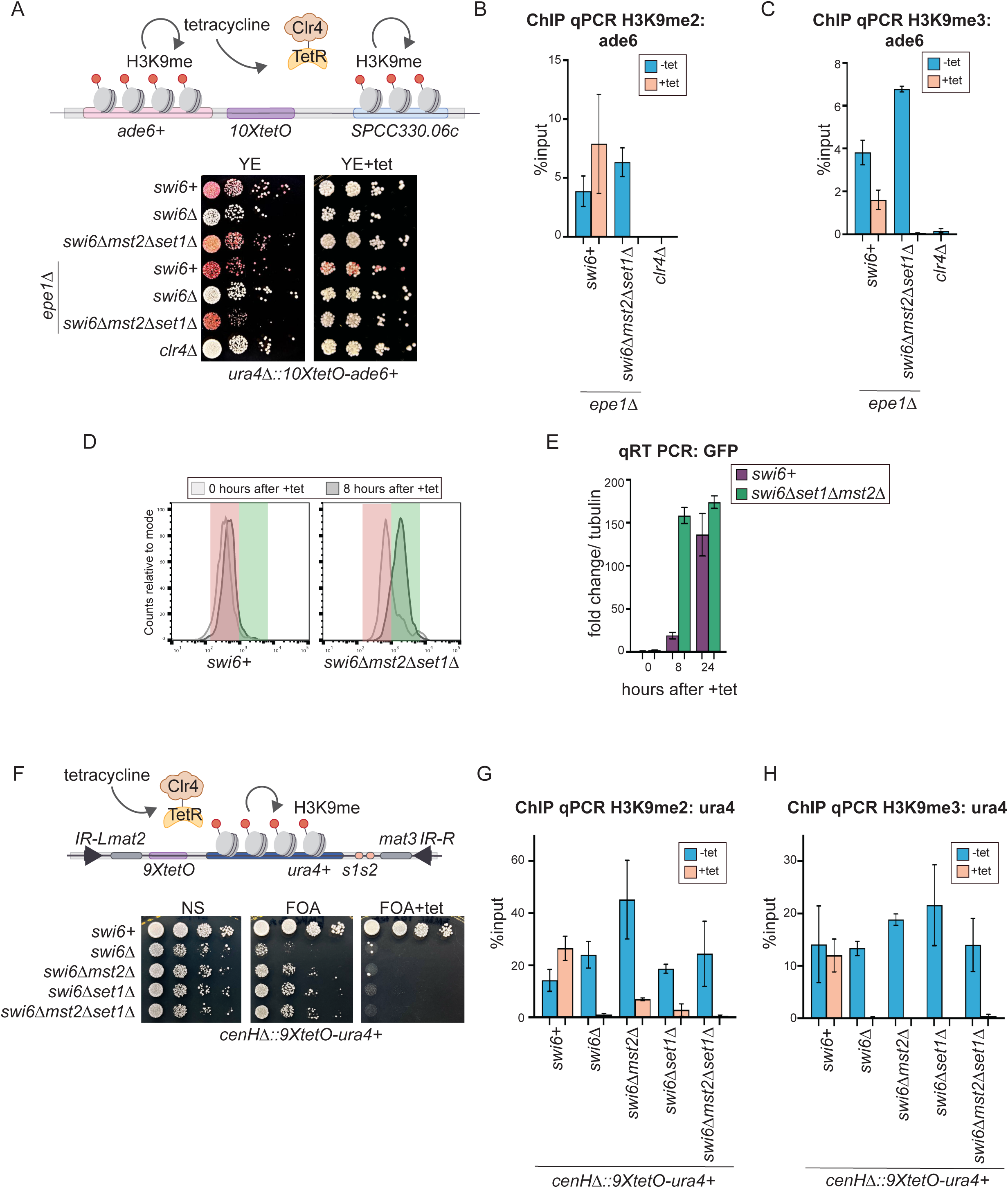
Swi6 is essential for heterochromatin maintenance even when Set1 and Mst2 are deleted. (**A**) Schematic of heterochromatin maintenance assay at the ectopic locus. Silencing assay of *ura4Δ::10XtetO*-*ade6+* reporter in indicated genotypes without or with tetracycline. (**B, C**) ChIP-qPCR of H3K9me2 (B) or H3K9me3 (C) before and after tetracycline addition. Error bars indicate SD (N=2). (**D**) Flow cytometry to examine GFP expression in *swi6+* and *swi6Δmst2Δset1Δ* cells at 0 hours and 8 hours after tetracycline addition. Histograms shifted to the left indicate a GFP-OFF population (red) and those shifted to the right indicate a GFP-ON population (green). (**E**) qRT-PCR measurements of RNA levels of *GFP* in *swi6+* and *swi6Δmst2Δset1Δ* cells at 0, 8, and 24 hours after tetracycline addition. Error bars indicate SD (N=3). (**F**) Schematic of heterochromatin maintenance assay at the mat locus. Silencing assay of *ura4+* reporter in the indicated genotypes with or without tetracycline. (**G, H**) ChIP-qPCR experiments to examine H3K9me2 (E) and H3K9me3 (F) before and after +tetracycline addition. Error bars indicate SD (N=2).

To quantitatively measure the dynamics of heterochromatin loss in *swi6Δmst2Δset1Δ* cells, we used a GFP reporter that enabled us to continuously track the loss of silencing after addition of tetracycline. After 8 hours of tetracycline addition which spans at least two cell divisions, *swi6Δmst2Δset1Δ* cells completely switched from a GFP-OFF state to a GFP-ON state unlike *swi6+* cells which successfully maintain the GFP-OFF state over this short time-period (**Figure 5D-E**).

Additionally, we used strains where TetR-Clr4-I is required for heterochromatin establishment at the *mat* locus.^46^ Plating *swi6+* cells on +tet containing media does not disrupt silencing as shown by the robust growth of cells on FOA+tet (**Figure 5F**, *swi6+*). Hence, *swi6+* cells can successfully maintain epigenetic silencing. In contrast, *swi6Δmst2Δset1Δ* cells that could successfully silence the *ura4+* reporter when TetR-Clr4-I was bound (despite the absence of Swi6) (**Figure 5F**), failed to maintain silencing on tetracycline containing media. *swi6Δmst2Δset1Δ* cells exhibited a near-complete loss of growth on FOA+tet. We also confirmed through ChIP-qPCR the loss of H3K9me2 and H3K9me3 at the *ura4+* reporter in *swi6Δmst2Δset1Δ* cells although this modification was successfully maintained in the presence of *swi6+* (**Figure 5G-H**). Hence, our results suggest that Swi6 may have a genetically separable function in maintaining epigenetic memory.

### Swi6 oligomerization is essential for epigenetic inheritance of H3K9 methylation

Previous studies have shown that HP1 proteins (including Swi6) oligomerize to form condensates that have liquid or gel-like properties.^19–21^ A mutation in the chromodomain of Swi6 (Swi6-LoopX, R93A K94A) attenuates Swi6 oligomerization by disrupting CD-CD interactions, which affects phase separation properties *in vitro* (**Figure 6A**).^14^ We replaced the wild-type copy of Swi6 with a Swi6-LoopX mutant in strains that have a *10XtetO-ade6+* reporter and an Epe1 deletion (*epe1Δ)*. In *swi6-LoopX* mutants, we initially observed red colonies on media lacking tetracycline and red or sectored colonies on media containing +tetracycline (**Figure 6B**). This phenotype resembled what we observed in cells that express wild-type Swi6. However, when cells were pre-grown in +tetracycline medium prior to plating, we observed that Swi6-LoopX expressing cells turned white. In contrast, cells expressing a wild-type copy of Swi6 robustly maintained silencing exhibiting red or sectored colonies. We confirmed through ChIP-qPCR measurements that H3K9me2 and H3K9me3 levels are comparable in cells with a wild-type copy of Swi6 or Swi6-LoopX during establishment (-tet, **Figure 6C-D**). These results suggest that Swi6 oligomerization is not required to establish epigenetic silencing at the ectopic locus. In contrast, ChIP-qPCR measurements of Swi6-LoopX cells (+tet, **Figure 6C-D**) showed a partial loss of H3K9me2 and a complete loss of H3K9me3 during maintenance. When we replaced Swi6 with Swi6-LoopX in strains with inducible heterochromatin at the *mat* locus, *swi6-LoopX* establishment and maintenance were unaffected compared to *swi6+*, as indicated by growth on FOA and FOA+tet medium (**Figure S4A**). Unlike the ectopic locus, the *mat* locus has additional DNA binding sequences and boundary elements, which may ensure that maintenance is robust in cells expressing the Swi6-LoopX mutant. It should be noted that the essential role of Swi6 in maintenance is still apparent at the *mat* locus in a *swi6Δ* or *swi6Δmst2Δset1Δ* background.

**Figure 6.**
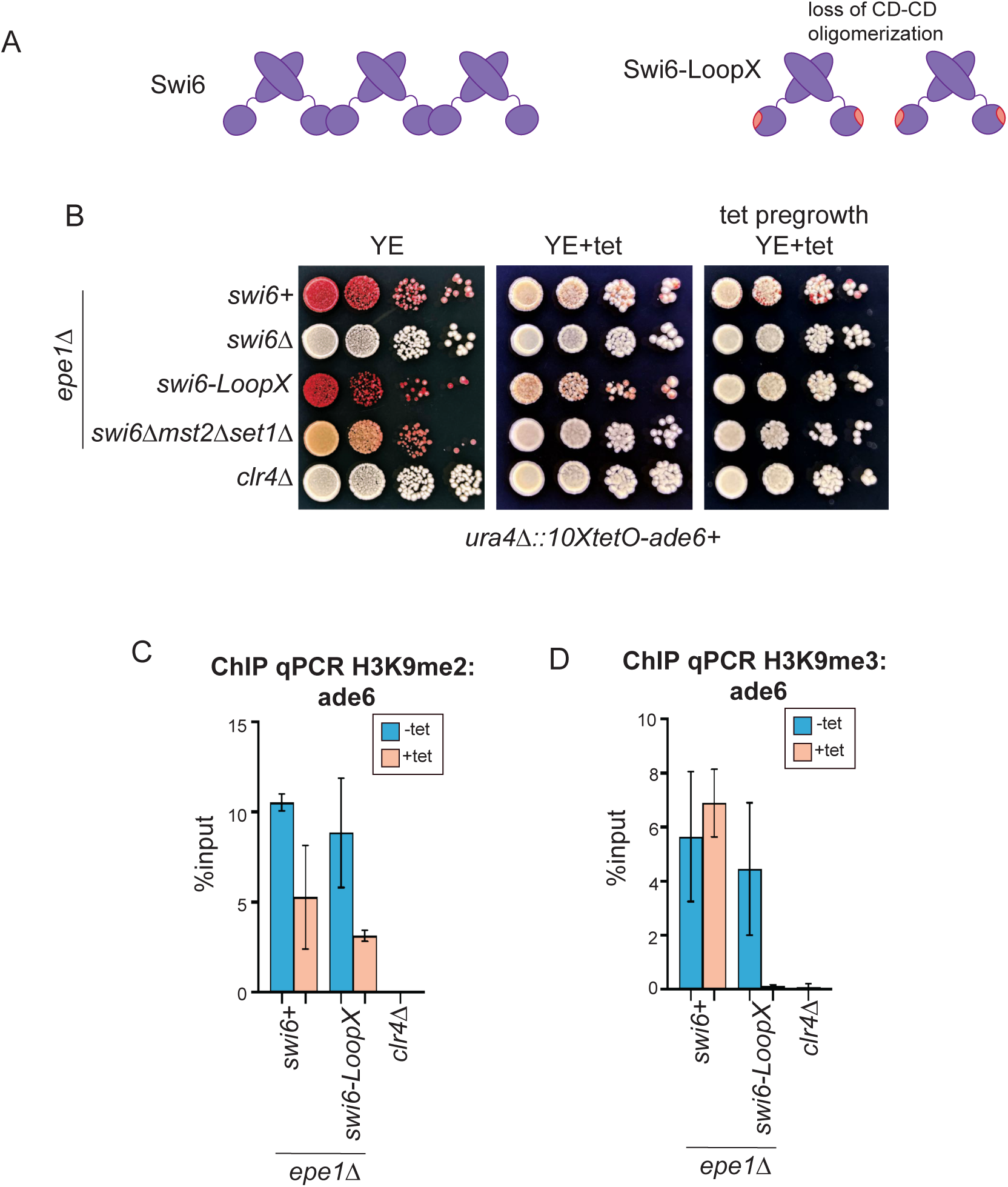
Swi6 oligomerization is required for maintenance. (**A**) Schematic of Swi6 oligomerization. Wild type Swi6 oligomerizes through CD-CD interactions. Swi6-LoopX (Swi6-R93A/K94A) disrupts Swi6 oligomerization. (**B**) Silencing assay of *ura4Δ::10XtetO*-*ade6+* reporter strains in the indicated genotypes. (**C,D**) ChIP-qPCR of H3K9me2 (**C**) or H3K9me3 (**D**) with +tetracycline to measure maintenance at the *ade6+* reporter. Error bars indicate SD (N=2). See also Figure S4.

## DISCUSSION

HP1 proteins play multifaceted roles in heterochromatin formation and epigenetic inheritance. These roles include promoting heterochromatin specific protein-protein interactions, chromatin compaction, and the spatial relocalization of heterochromatin domains to the nuclear periphery.^9,10,18,67^ Here, we investigate the function of the major *S. pombe* HP1 protein, Swi6, and show that it has genetically separable roles in heterochromatin initiation, establishment, and maintenance (**Figure 7**). Our data suggests that Swi6 contributes to heterochromatin establishment through protein-protein interactions, rather than its inherent structural or biophysical properties. In addition, we discovered that deleting Set1 and Mst2 can also lead to the bypass of Chp2 (the second *S.pombe* HP1 protein) when Epe1, the primary H3K9 demethylase like enzyme in *S.pombe,* is absent (**Figure S1E**). These results point to different genetic requirements for the bypass of the two *S.pombe* HP1 proteins given their non-overlapping functions in heterochromatin formation.^29^ Importantly, our findings imply that HP1-mediated chromatin compaction or protein exclusion due to HP1-mediated phase separation is unlikely to be the molecular basis for heterochromatin silencing.^18,19,21,68^

**Figure 7.**
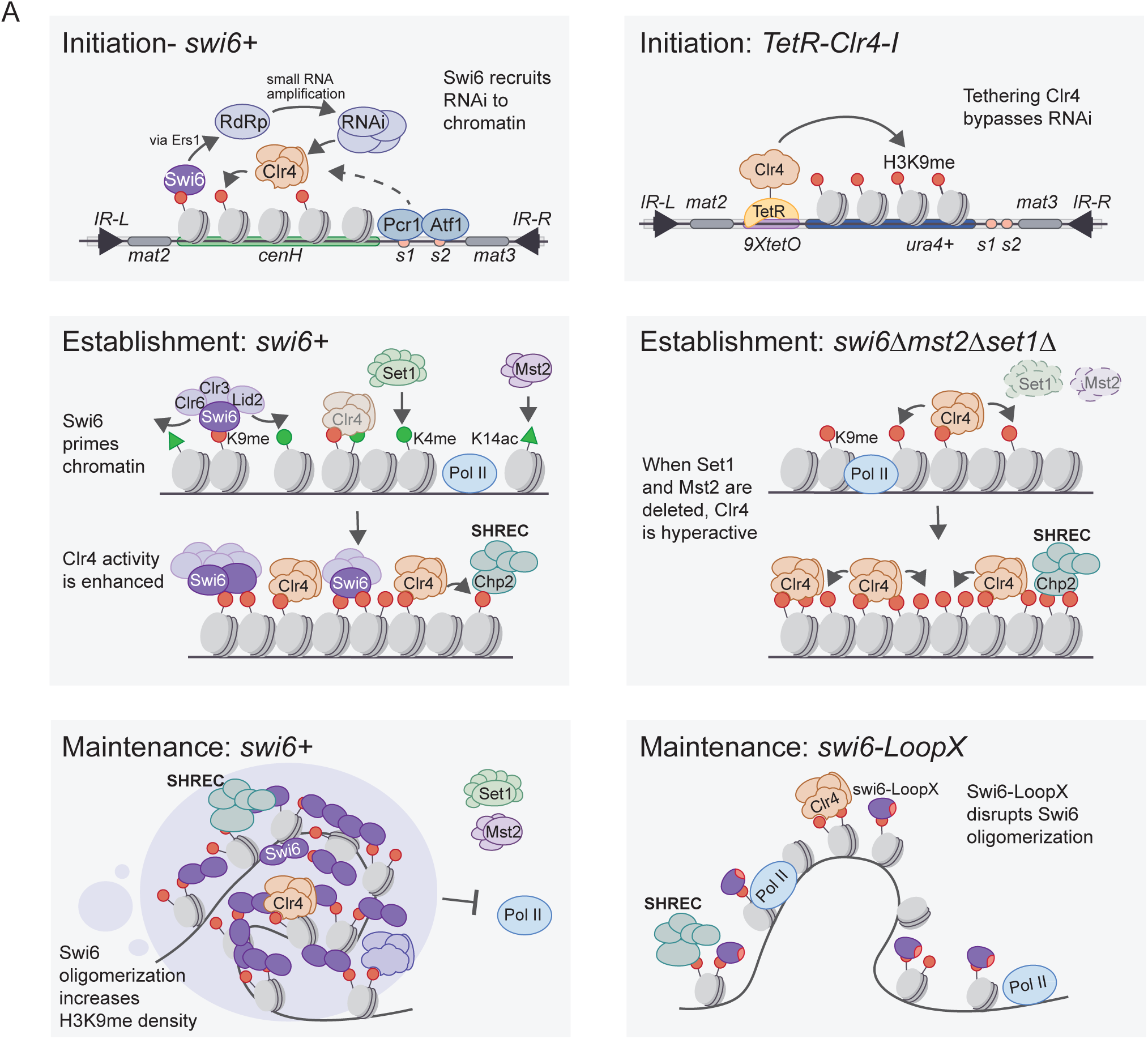
Model: Swi6 can be bypassed during heterochromatin establishment but is indispensable during heterochromatin maintenance. Initiation: Swi6 is required for initiation at RNAi dependent sites of heterochromatin formation. This process can be bypassed by Clr4 tethering. Establishment: The initial recruitment of Clr4 to specific sites in the genome leads to low levels of H3K9me. This is followed by Swi6 binding to recruit histone modifiers that prime chromatin which in turn enhances Clr4 enzymatic activity. Swi6 priming involves the removal of H3K4me and H3K14ac both of which are modifications catalyzed by proteins associated with active transcription-Set1 and Mst2. Swi6 can be bypassed when Set1 and Mst2 are deleted. Maintenance: Swi6 binds to and oligomerizes in complex with H3K9me chromatin. These structural features of Swi6 binding to heterochromatin promote long-range interactions that enhance H3K9me density leading to robust epigenetic inheritance. Maintenance is lost when Swi6 oligomerization is disrupted in Swi6-LoopX mutants.

At the pericentromeric repeats and the mating type locus, Swi6 binds Ers1, which acts as an adaptor to recruit the RNA-dependent RNA polymerase complex (RDRC) involved in double-stranded RNA synthesis.^31–33^ In this study, we bypassed the requirement for Swi6 in initiation via TetR-Clr4-I recruitment, which enabled us to investigate the consequences of HP1 function downstream of H3K9 methylation. Our results underscore the importance of using site-specific tethering and inducible systems to investigate complex, overlapping chromatin interactions *in vivo*.^69^

In our experiments, we found that the deletion of Set1 and Mst2 renders Swi6 dispensable for silencing at both an ectopic site and the endogenous mating type locus. HP1 proteins lack catalytic functions, so Swi6 likely recruits histone-modifying enzymes such as Lid2, an H3K4 demethylase, Clr3, an H3K14 deacetylase, and Clr6, an H3K9 deacetylase.^10^ Our proposed stepwise model of heterochromatin establishment involves an initial phase of Clr4 recruitment, which leads to low levels of H3K9me (**Figure 7**). Swi6 then directs the recruitment of various histone modifiers, priming chromatin to enhance Clr4 activity further.

Previous studies have shown silencing can occur independently of Swi6 in a synthetic context where Clr4 lacking its chromodomain (Clr4ΔCD) is recruited to an ectopic site using a Gal4 fusion protein.^70^ This observation suggests that the role of Swi6 is limited to recruiting RNAi factors to pericentromeric repeats, making it dispensable in the context of an RNAi-independent heterochromatin initiation system. However, other studies have shown that Swi6 has a crucial role in transcriptional gene silencing, a downstream step of H3K9 methylation deposition.^29,57^ Our system, in which we express both the TetR-Clr4DCD initiator and a full-length copy of Clr4, recapitulates the essential role of Swi6 in transcriptional gene silencing (**Figure 1C-D**).

A canonical role ascribed to HP1 proteins is heterochromatin spreading wherein H3K9 methylation extends across large chromosomal domains to sites that are distal from nucleation elements (∼10-100s of kilobases).^17^ In the contexts where we observed Swi6 independent silencing, we could also completely restore H3K9 methylation in excess of what is present in wild-type cells. Therefore, we favor a model in which the activity of Clr4 is the primary determinant of H3K9 methylation spreading instead of requiring processive HP1-dependent mechanisms.

Recent work has highlighted that HP1 proteins form condensates that have liquid or gel-like properties *in vitro* and in cells.^19–21^ Whether the material properties of HP1-chromatin complexes are most consistent with phase separation remains contested.^71^ Regardless, both condensate-dependent and independent models require HP1 oligomerization. Indeed, we have demonstrated that Swi6 oligomerization is not required for heterochromatin establishment in two strain backgrounds, *swi6Δmst2Δset1Δ* and *swi6-LoopX* (**Figure 6B**).^14^ Furthermore, RNA polymerase II occupancy in *swi6Δmst2Δset1Δ* cells is equal to that of *swi6+* cells suggesting that mechanisms other than condensate-based exclusion can also lead to the same degree of transcriptional gene silencing (**Figure 1G**).

Deleting Swi6 or expressing the Swi6 LoopX mutant selectively disrupts epigenetic inheritance (**Figure 6A**). Oligomerization could promote various functions, including bridging, compaction, or phase separation, that are important only during maintenance. For example, HP1 phase separated condensates, could selectively include chromatin factors that may have a maintenance-specific function. Heterochromatin is important for long range chromatin contacts and increases chromatin compaction.^72^ Therefore, HP1 oligomerization independent of phase separation could mediate long-range chromosomal interactions or promote chromatin compaction which may be essential only during maintenance. These processes may ultimately lead to an increase in the density of H3K9me which is important for epigenetic inheritance.^65^ Hence, our data suggests that HP1 proteins may be essential to create H3K9me enriched chromatin networks that facilitate the re-recruitment of Clr4 to propagate stable epigenetic states.

In summary, our studies show that HP1 proteins, primarily function as adapters that can be repurposed to recruit diverse histone modifying proteins during establishment. Our results may explain the diversification of HP1 proteins as transcriptional activators since this would simply require the evolution of new protein-protein interactions or as cross-linkers that preserve the mechanical stability of the nucleus. In addition, HP1 proteins appear to have a genetically separable structural role during maintenance. These observations are in part consistent with recent mathematical models of how proteins that affect 3-D genome structure (such as HP1 proteins as shown in our study) in combination with limiting amounts of the enzyme are essential features to build chromatin-based systems that encode epigenetic memory.^73,74^

## LIMITATIONS

Our study clearly establishes a genetic basis for the bypass of the major HP1 protein, Swi6 during heterochromatin establishment. Although hypermethylation arising from deleting Set1 and Mst2 can serve as one bypass mechanism, it is likely that more extensive genetic screens may reveal new and unexpected chromatin features that additionally contribute to this process. Since Swi6 recruits the major heterochromatin antagonist Epe1, loss of Epe1 recruitment could additionally contribute to the bypass mechanisms we observed in this study. Our studies define what is minimally required for epigenetic silencing by ruling out HP1 dependent compaction or HP1 dependent condensate-based models of heterochromatin formation. Yet, our studies are based on genetic evidence and lack the precision that *in vitro* reconstitution methods can afford to assess the roles of HP1 proteins more directly in this process. It is possible that condensates can form within deacetylated heterochromatin domains in an HP1 dependent manner which could effectively have critical roles in establishing silencing. We have identified a Swi6 separation of function mutant where disrupting Swi6 oligomerization selectively disrupts epigenetic inheritance. It is unclear whether disrupting oligomerization affects Swi6 alone or if these mutations may also affect protein-protein interactions. Finally, Swi6 is a highly dynamic protein that turns over from sites of heterochromatin formation on the millsecond to second timescale. How transient Swi6 binding leads to stable and heritable chromatin states is not explained by our data.

## ACKNOWLEDGMENTS

The authors declare no competing interests. We thank Danesh Moazed for sharing strains used in this study. We thank Magdalena Murwaska and Sigurd Braun for sharing strains for RITE assays. We thank Nidhi Khurana and Gulzhan Raiymbek for their support in obtaining preliminary data during the initial phase of this study and Saikat Biswas for help with running samples on the flow cytometer. We thank Amanda Ames and Basila Moochickal Assainar for their supportive and insightful feedback regarding this study. This work was funded by NIH award R35GM137832 to KR, T32GM007315 to MS, and T32GM007544 to AL.

## AUTHOR CONTRIBUTIONS

Conceptualization, M.S. and K.R.; Methodology, M.S. and K.R.; Formal Analysis, M.S and A.J.L.; Investigation, M.S., A.L., and F.H.; Writing – Original Draft, M.S. and K.R.; Writing – Review & Editing, M.S. and K.R.; Funding Acquisition, M.S., A.J.L., and K.R.; Supervision, M.S. and K.R.

## INCLUSION AND DIVERSITY

We support inclusive, diverse, and equitable conduct of research.

## STAR★Methods

### Key resources table

| REAGENT or RESOURCE | SOURCE | IDENTIFIER |
| --- | --- | --- |
| Antibodies |  |  |
| anti-H3K9me2 | Abcam | ab1220 |
| anti-H3K9me3 | Active Motif | Cat No. 39161 |
| anti-H3K9me3 | Diagenode | Cat No. C15410193 |
| anti-H3 | Abcam | ab1791 |
| anti-H3K4me3 | Abcam | ab8580 |
| anti-H3K14ac | Abcam | ab52946 |
| anti-Rpb1 | Cell Signaling | Cat No. 2629 |
| anti-T7 | Merck | 69522–3 |
| Chemicals, Peptides, and Recombinant Proteins |  |  |
| 5-Fluorootic acid monohydrate (FOA) | GoldBio | F-230 |
| Dynabeads Protein A | Invitrogen | 10002D |
| Formaldehyde | Electron Microscopy Sciences | 15680 |
| Glycogen | Thermo Scientific | R0561 |
| Hygromycin B | GoldBio | H-270 |
| Proteinase K | Invitrogen | 25530049 |
| Phenol chloroform isoamyl alcohol | Sigma-Aldrich | 77617 |
| Anhydrous Tetracycline (tet) | Cayman Chemical | 10009542100 |
| G-418 Sulfate | GoldBio | G-418 |
| Mag-Bind Total Pure NGS | Omega | M1378 |
| PVDF Blotting Membrane | GE Healthcare | 10600022 |
| EMM powder | Formedium | PMD0402 |
| Yeast extract | Gibco | 2911929 |
| DNase I | NEB | M0303 |
| Phenol chloroform, acidic | VWR | E277 |
| Adenine hemisulfate salt | Chem-Impex Int'l | 00006 |
| Streptavidin magnetic beads | NEB | S1420 |
| Dimethyl adipimidate dihydrochloride | Sigma Aldrich | 285625 |
| Hydroxyurea | EMD Millipore Corp | 400046 |
| β-estradiol | MP Biomedicals | 194565 |
| Critical Commercial Assays |  |  |
| Superscript III Reverse Transcriptase | Invitrogen | 18080085 |
| Syber Green | BioRad | 172-5120 |
| NEBNext Ultra II FS DNA Library Prep Kit for Illumina | NEB | E7805 |
| NEBNext Multiplex Oligos | NEB | E6609 |
| Qubit dsDNA HS assay kit | Thermo Fisher | Q32854 |
| RNeasy Mini Kit | Qiagen | 74104 |
| Deposited Data |  |  |
| H3K9me2/H3K9me3 ChIP-seq | This study | GSE244017 |
| RNA-seq | This study | GSE233333 |
| Experimental Models: Organisms/Strains |  |  |
| *S. pombe* | This study | Table S1 |
| Oligonucleotides |  |  |
| qRT-PCR, ChIP-qPCR | This study | Table S2 |
| Software and Algorithms |  |  |
| Prism v9.4.1 | GraphPad Software Inc. | https://www.graphpad.com/scientific-software/prism/ |
| FlowJo v10.9.0 | Becton, Dickinson and Company |  |
| bwa v0.7.17 - r1188 | Wellcome Trust | http://bio-bwa.sourceforge.net |
|  | Sanger Institute |  |
| R (version 4.0.2) | Cran.R | https://cran.r-project.org/ |
| ggplot2 | Cran.R | https://ggplot2.tidyverse.org/ |
| Trimmomatic v 0.39 | Max Planck Institute of Molecular Plant Physiology | http://www.usadellab.org/cms/?page=trimmomatic |
| RStudio (version 1.4.1106) | RStudio | https://www.rstudio.com/ |
| USeq v9.2.9 | Utah Bioinformatics Shared Resource Center | https://useq.sourceforge.net |
| deeptools v3.5.1 | Max Planck Institute | https://github.com/deeptools/deepTools/blob/master/docs/content/about.rst |
| IGV v11.0.13 | Broad Institute and the Regents of the University of California | https://software.broadinstitute.org/software/igv/ |
| MACS2 |  | https://github.com/macs3-project/MACS |
| samtools v1.6 | Wellcome Trust Sanger Institute | http://samtools.sourceforge.net/ |
| bedtools v2.27.1 | University of Utah | https://bedtools.readthedocs.io/en/latest/ |
| STAR |  | https://github.com/alexdobin/STAR |
| SnapGene v5.0.8 |  | www.snapgene.com |

### Resource availability

#### Lead Contact

Further information and requests for resources and reagents should be directed to and will be fulfilled by the lead contact, Kaushik Ragunathan.

#### Materials Availability

All unique/stable reagents generated in this study are available from the lead contact without restriction.

#### Data and Code Availability

- ChIP-seq and RNA-seq data have been deposited at GEO and are publicly available as of the date of publication. ChIP-Seq accession number is GSE244017. RNA-Seq accession number is GSE233333.
- This paper does not report original code.
- Any additional information required to reanalyze the data reported in this paper is available from the lead contact upon request.

#### Experimental model and study participant details

All *S. pombe* strains in this study are listed in Table S1. Some strains were made using a PCR-based gene targeting approach or with CRISPR to insert point mutations or increase transformation efficiency ^75,76^. Other strains were made by a cross followed by random spore analysis. Strains with point mutations were confirmed with sequencing. Strains were grown at 32C in YEA media.

### Method details

#### Western blot

A standard TCA precipitation protocol was used to extract protein from 5 OD of cells. Briefly, pellets were washed with 1ml of ice-cold water, then resuspended in 150ul of YEX buffer (1.85M NaOH, 7.5% beta-mercaptoethanol) and incubated on ice for ten minutes. Then 150ul of 50% TCA was mixed into each sample and incubated on ice for ten minutes. Samples were then spun down for five minutes at 13000rpm at 4C. The pellets were resuspended in 40ul of SDS sample buffer (125mM Tris buffer pH 6.8, 8M urea, 5% SDS, 20% glycerol, 5% BME) and spun down for five minutes at 13000rpm at 4C. 10ul was then run on a gradient (4-20%) SDS-PAGE gel at 200V. Gel transfer was performed on a Trans-Blot Turbo Transfer to a PVDF membrane. The membrane was blocked for 45 min with 5% milk in TBS pH7.5 with 0.1% Tween-20 (TBST). The membrane was incubated overnight with primary antibody (H3 (Abcam, ab1791), H3K4me (Abcam, ab8580), or H3K14ac(Abcam, ab52946)) at 4C. Membranes were washed three times with TBST and incubated with secondary antibody for one hour at RT. Membranes were then incubated with chemiluminescence substrate and imaged using a BioRad ChemiDoc imager.

#### Chromatin immunoprecipitation (ChIP)

30 ml of cells were grown to late log phase (OD_600_- 1.8-2.2) in yeast extract supplemented with adenine or yeast extract supplemented with adenine containing tetracycline (2.5µg/ml) and fixed with 1% formaldehyde for 15 min at room temperature (RT). For Rpb1 samples, cells were resuspended in PBS with 0.25% DMSO and crosslinked with 10 mM DMA for 45 minutes at RT. Then washed once with PBS and resuspended in PBS to crosslink with 1% formaldehyde for 15min at RT. 130mM glycine was then added to quench the reaction and incubated for 5 min at RT. The cells were harvested by centrifugation, and washed with TBS (50 mM Tris, pH 7.6, 500 mM NaCl). Cell pellets were resuspended in 300 µl lysis buffer (50 mM HEPES-KOH, pH 7.5, 100 mM NaCl, 1mM EDTA, 1% Triton X-100, 0.1% SDS, and protease inhibitors) to which 500 µL 0.5 mm glass beads were added and cell lysis was carried out by bead beating using Omni Bead Ruptor at 3000 rpm x 30sec x 10 cycles. Tubes were punctured and the flow-through was collected in a new tube by centrifugation which was subjected to sonication to obtain fragment sizes of roughly 100-500 bp long. After sonication, the extract was centrifuged for 15 min at 13000 rpm at 4°C. The soluble chromatin was then transferred to a fresh tube and normalized for protein concentration by the Bradford assay. For each normalized sample, 25µL lysate was saved as input, to which 225 µL of 1xTE/1% SDS were added (TE: 50 mM Tris pH 8.0, 1 mM EDTA). Protein A Dynabeads were preincubated with antibody For the H3K9me3 antibody (C15410193, Diagenode) preincubation was done with Streptavidin beads. For each immunoprecipitation, 2 µg H3K9me2 antibody (ab1220, Abcam), 2µg H3K9me3 antibody (39161, Active Motif), 4 µg H3K9me3 antibody (C15410193, Diagenode), or 2µg T7 antibody (69522–3, Merck) coupled to 30µL beads was added to 400µL soluble chromatin (Diagenode H3K9me3 antibody was used in Figure 2E only) and the final volume of 500µL was achieved by adding lysis buffer. Samples were incubated for 3h at 4°C. The beads were collected on magnetic stands and washed 3 times with 1 ml lysis buffer and once with 1 ml TE. For eluting bound chromatin, 100µL elution buffer I (50 mM Tris pH 8.0, 10mM EDTA, 1% SDS) was added, and the samples were incubated at 65°C for 5 min. The eluate was collected and incubated with 150 µL 1xTE/0.67% SDS in the same way. Input and immunoprecipitated samples were finally incubated overnight at 65°C to reverse crosslink. 60µg glycogen, 100 µg proteinase K (Roche), 44 ul of 5M LiCl, and 250 ul of 1xTE was added to each sample, and incubation was continued at 55°C for 1h. Phenol/chloroform extraction was carried out for all the samples, followed by ethanol precipitation. Immuno-precipitated DNA was resuspended in 100µL of 10 mM Tris pH 7.5 and 50 mM NaCl. ChIP experiments were analyzed using quantitative PCR with Taq polymerase and SYBR Green using a CFX Opus 384 Real-Time PCR System. All data is presented as percent enrichment relative to input.^77^

#### ChIP-Seq library preparation and processing

Libraries were constructed using the manufacturer’s guidelines in the NEBNext® Ultra™ II FS DNA Library Prep Kit for Illumina, using 1ng of starting material. Barcoded libraries were pooled and sequenced with next-generation sequencing. First, raw reads were demultiplexed by barcode. Then the sequences were trimmed with Trimmomatic, aligned with BWA, and normalized by reads per million.^78,79^ Then the reads were visualized with IGV. For further analysis peaks were called using MACS2 with -g 12.57e6 in broad mode with a cutoff of 0.05.^80^ Heatmaps were generated using deepTools (v3.5.1).^81^

#### Histone turnover assay

Cells were inoculated in YEA supplemented with Hygromycin B (50mg/mL) and grown O/N at 32C to an OD of 0.2-0.4.10 ODs of cells were harvested for ChIP as the 0h time point. 10 ODs of the remaining cells were washed twice with YEA without Hygromycin B. Then resuspended in YEA with 15mM Hydroxyurea (HU) and 1.5mM β -estradiol (ER). After four hours of growth at 32C, 10 ODs of cells were harvested for ChIP as the 4 h time point.

#### RNA extraction

10ml of cells were grown to late log phase (OD_600_- 1.8-2.2) in yeast extract supplemented with adenine. Cells were resuspended in 750uL TES buffer (0.01M Tris pH7.5, 0.01M EDTA, 0.5% SDS). Immediately 750uL of acidic phenol chloroform was added and vortexed for 2 minutes. Samples were incubated at 65C for forty minutes, vortexing for 20 seconds every ten minutes. The aqueous phase was separated by centrifuging in Phase Lock tubes for 5 minutes at 13000 rpm at 4C. The aqueous phase was transferred to new tubes and ethanol precipitated. After extraction, RNA was treated with DNase. Then the RNA was cleaned up using RNeasy Mini kits (QIAGEN). cDNA was prepared using oligoDT and SuperScript III Reverse Transcriptase (Invitrogen). The cDNA was then used for qPCR with SYBR Green and Taq polymerase on a CFX OPUS 384 Real-Time PCR System. Relative RNA is quantified using ΔCT compared to *tub* levels.

#### RNA-seq analysis

Libraries were prepared, and sequencing was performed commercially. Raw fastq files were aligned to the ASM294v2 reference genome using STAR and then indexed using samtools.^82^ Bam files were grouped by genotype replicate and differential expression analysis was performed through Defined Region Differential Seq in the USEQ program suite.^83^ After Benjamini and Hochberg multiple testing corrections, the cutoff for significant differential expression of pairwise gene comparisons was defined as a P value of <0.01 (prior to phred transformation). Volcano plots were drawn using the ggplot2 library, and heatmap was drawn using the pheatmap library, as well as the standard R library and functions.

#### Silencing assays

Cells were grown in 3 ml of yeast extract containing adenine (YEA) at 32°C overnight. Cells were washed twice in water and then resuspended to a concentration of 3 x 10^7^ cells/ml. 5µl of ten-fold serial dilutions was spotted on indicated plates (YE, YE+tet, EMMC, EMMC+FOA, EMMC+FOA+tet). Plates were incubated for 2-5 days before photographing.

#### FACS analysis

Cells containing TetR-Clr4 and a *ura4-GFP* reporter were harvested in log phase. 3 x 10^7^ cells/ml were washed with water and then fixed with 70% ethanol/30% sorbitol. The cells were washed three times with 1X tris-buffered saline (TBS) and resuspended in 1ml 1xTBS. GFP fluorescence was measured using a Bio-Rad ZE5 cell analyzer with wavelength 488nm. A primary gate based on physical parameters (forward and side light scatter) was set to exclude dead cells or debris. 10,000 cells were analyzed for each sample.

### Quantification and statistical analysis

Prism was used to make all graphs and quantify data. Number of replicates is indicated in each figure legend. Error bars of qRT-PCR and ChIP-qPCR represent standard deviation.

## REFERENCES

1. Luger, K., Mader, A.W., Richmond, R.K., Sargent, D.F., and Richmond, T.J. (1997). Crystal structure of the nucleosome core particle at 2.8 Å resolution. Nature 389, 251–260. 10.1038/38444.

2. Li, B., Carey, M., and Workman, J.L. (2007). The Role of Chromatin during Transcription. Cell 128, 707–719. 10.1016/j.cell.2007.01.015.

3. Kouzarides, T. (2007). Chromatin Modifications and Their Function. Cell 128, 693–705. 10.1016/j.cell.2007.02.005.

4. Allshire, R.C., and Madhani, H.D. (2018). Ten principles of heterochromatin formation and function. Nat. Rev. Mol. Cell Biol. 19, 229–244. 10.1038/nrm.2017.119.

5. Bannister, A.J., Zegerman, P., Partridge, J.F., Miska, E.A., Thomas, J.O., Allshire, R.C., and Kouzarides, T. (2001). Selective recognition of methylated lysine 9 on histone H3 by the HP1 chromo domain. Nature 410, 120–124. 10.1038/35065138.

6. Lachner, M., O’Carroll, D., Rea, S., Mechtler, K., and Jenuwein, T. (2001). Methylation of histone H3 lysine 9 creates a binding site for HP1 proteins. Nature 410, 116–120. 10.1038/35065132.

7. Cowieson, N.P., Partridge, J.F., Allshire, R.C., and McLaughlin, P.J. (2000). Dimerisation of a chromo shadow domain and distinctions from the chromodomain as revealed by structural analysis. Curr. Biol. 10, 517–525. 10.1016/S0960-9822(00)00467-X.

8. Jacobs, S.A., and Khorasanizadeh, S. (2002). Structure of HP1 Chromodomain Bound to a Lysine 9-Methylated Histone H3 Tail. Science 295, 2080–2083. 10.1126/science.1069473.

9. Thiru, A., Nietlispach, D., Mott, H.R., Okuwaki, M., Lyon, D., Nielsen, P.R., Hirshberg, M., Verreault, A., Murzina, N.V., and Laue, E.D. (2004). Structural basis of HP1/PXVXL motif peptide interactions and HP1 localisation to heterochromatin. EMBO J. 23, 489–499. 10.1038/sj.emboj.7600088.

10. Fischer, T., Cui, B., Dhakshnamoorthy, J., Zhou, M., Rubin, C., Zofall, M., Veenstra, T.D., and Grewal, S.I.S. (2009). Diverse roles of HP1 proteins in heterochromatin assembly and functions in fission yeast. Proc. Natl. Acad. Sci. 106, 8998–9003. 10.1073/pnas.0813063106.

11. Eissenberg, J.C., and Elgin, S.C.R. (2014). HP1a: a structural chromosomal protein regulating transcription. Trends Genet. 30, 103–110. 10.1016/j.tig.2014.01.002.

12. Machida, S., Takizawa, Y., Ishimaru, M., Sugita, Y., Sekine, S., Nakayama, J. ichi, Wolf, M., and Kurumizaka, H. (2018). Structural Basis of Heterochromatin Formation by Human HP1. Mol. Cell 69, 385–397.e8. 10.1016/j.molcel.2017.12.011.

13. Canzio, D., Chang, E.Y., Shankar, S., Kuchenbecker, K.M., Simon, M.D., Madhani, H.D., Narlikar, G.J., and Al-Sady, B. (2011). Chromodomain-mediated oligomerization of HP1 suggests a nucleosome-bridging mechanism for heterochromatin assembly. Mol. Cell 41, 67–81. 10.1016/j.molcel.2010.12.016.

14. Canzio, D., Liao, M., Naber, N., Pate, E., Larson, A., Wu, S., Marina, D.B., Garcia, J.F., Madhani, H.D., Cooke, R., et al. (2013). A conformational switch in HP1 releases auto-inhibition to drive heterochromatin assembly. Nature 496, 377–381. 10.1038/nature12032.

15. Hinde, E., Cardarelli, F., and Gratton, E. (2015). Spatiotemporal regulation of Heterochromatin Protein 1-alpha oligomerization and dynamics in live cells. Sci. Rep., 12001. 10.1038/srep12001.

16. Schmiedeberg, L., Weisshart, K., Diekmann, S., Meyer zu Hoerste, G., and Hemmerich, P. (2004). High- and Low-mobility Populations of HP1 in Heterochromatin of Mammalian Cells. Mol. Biol. Cell 15, 2819–2833.

17. Verschure, P.J., van der Kraan, I., de Leeuw, W., van der Vlag, J., Carpenter, A.E., Belmont, A.S., and van Driel, R. (2005). In Vivo HP1 Targeting Causes Large-Scale Chromatin Condensation and Enhanced Histone Lysine Methylation. Mol. Cell. Biol. 25, 4552–4564. 10.1128/MCB.25.11.4552-4564.2005.

18. Keenen, M.M., Brown, D., Brennan, L.D., Renger, R., Khoo, H., Carlson, C.R., Huang, B., Grill, S.W., Narlikar, G.J., and Redding, S. (2021). HP1 proteins compact DNA into mechanically and positionally stable phase separated domains. eLife 10, e64563. 10.7554/eLife.64563.

19. Larson, A.G., Elnatan, D., Keenen, M.M., Trnka, M.J., Johnston, J.B., Burlingame, A.L., Agard, D.A., Redding, S., and Narlikar, G.J. (2017). Liquid droplet formation by HP1α suggests a role for phase separation in heterochromatin. Nature 547, 236–240. 10.1038/nature22822.

20. Strom, A.R., Emelyanov, A.V., Mir, M., Fyodorov, D.V., Darzacq, X., and Karpen, G.H. (2017). Phase separation drives heterochromatin domain formation. Nature 547, 241–245. 10.1038/nature22989.

21. Sanulli, S., Trnka, M.J., Dharmarajan, V., Tibble, R.W., Pascal, B.D., Burlingame, A.L., Griffin, P.R., Gross, J.D., and Narlikar, G.J. (2019). HP1 reshapes nucleosome core to promote phase separation of heterochromatin. Nature 575. 10.1038/s41586-019-1669-2.

22. Nishibuchi, G., and Nakayama, J. (2014). Biochemical and structural properties of heterochromatin protein 1: understanding its role in chromatin assembly. J. Biochem. (Tokyo) 156, 11–20. 10.1093/jb/mvu032.

23. Larson, A.G., and Narlikar, G.J. (2018). The Role of Phase Separation in Heterochromatin Formation, Function, and Regulation. Biochemistry 57, 2540–2548. 10.1021/acs.biochem.8b00401.

24. Kumar, A., and Kono, H. (2020). Heterochromatin protein 1 (HP1): interactions with itself and chromatin components. Biophys. Rev. 12, 387–400. 10.1007/s12551-020-00663-y.

25. Schoelz, J.M., and Riddle, N.C. (2022). Functions of HP1 proteins in transcriptional regulation. Epigenetics Chromatin 15, 14. 10.1186/s13072-022-00453-8.

26. Grewal, S.I.S., and Jia, S. (2007). Heterochromatin revisited. Nat. Rev. Genet. 8, 35–46. 10.1038/nrg2008.

27. Nakayama, J., Rice, J.C., Strahl, B.D., Allis, C.D., and Grewal, S.I.S. (2001). Role of histone H3 lysine 9 methylation in epigenetic control of heterochromatin assembly. Science 292, 110–113. 10.1126/science.1060118.

28. Sugiyama, T., Cam, H.P., Sugiyama, R., Noma, K. ichi, Zofall, M., Kobayashi, R., and Grewal, S.I.S. (2007). SHREC, an Effector Complex for Heterochromatic Transcriptional Silencing. Cell 128, 491–504. 10.1016/j.cell.2006.12.035.

29. Motamedi, M.R., Hong, E.J.E., Li, X., Gerber, S., Denison, C., Gygi, S., and Moazed, D. (2008). HP1 Proteins Form Distinct Complexes and Mediate Heterochromatic Gene Silencing by Nonoverlapping Mechanisms. Mol. Cell 32, 778–790. 10.1016/j.molcel.2008.10.026.

30. Halic, M., and Moazed, D. (2010). Dicer-Independent Primal RNAs Trigger RNAi and Heterochromatin Formation. Cell 140, 504–516. 10.1016/j.cell.2010.01.019.

31. Rougemaille, M., Shankar, S., Braun, S., Rowley, M., and Madhani, H.D. (2008). Ers1, a rapidly diverging protein essential for RNA interference-dependent heterochromatic silencing in Schizosaccharomyces pombe. J. Biol. Chem. 283, 25770–25773. 10.1074/jbc.C800140200.

32. Hayashi, A., Ishida, M., Kawaguchi, R., Urano, T., Murakami, Y., and Nakayama, J.I. (2012). Heterochromatin protein 1 homologue Swi6 acts in concert with Ers1 to regulate RNAi-directed heterochromatin assembly. PNAS 109, 6159–6164. 10.1073/pnas.1116972109.

33. Rougemaille, M., Braun, S., Coyle, S., Dumesic, P.A., Garcia, J.F., Isaac, R.S., Libri, D., Narlikar, G.J., and Madhani, H.D. (2012). Ers1 links HP1 to RNAi. Proc. Natl. Acad. Sci. U. S. A. 109, 11258–11263. 10.1073/pnas.1204947109.

34. Jia, S., Noma, K., and Grewal, S.I.S. (2004). RNAi-Independent Heterochromatin Nucleation by the Stress-Activated ATF/CREB Family Proteins. Science 304, 1971–1976. 10.1126/science.1099035.

35. Kanoh, J., Sadaie, M., Urano, T., and Ishikawa, F. (2005). Telomere Binding Protein Taz1 Establishes Swi6 Heterochromatin Independently of RNAi at Telomeres. Curr. Biol. 15, 1808–1819. 10.1016/j.cub.2005.09.041.

36. Partridge, J.F., Borgstrøm, B., and Allshire, R.C. (2000). Distinct protein interaction domains and protein spreading in a complex centromere. Genes Dev. 14, 783–791. 10.1101/gad.14.7.783.

37. Danzer, J.R., and Wallrath, L.L. (2004). Mechanisms of HP1-mediated gene silencing in *Drosophila*. Development 131, 3571–3580. 10.1242/dev.01223.

38. Yamada, T., Fischle, W., Sugiyama, T., David Allis, C., and Grewal, S.I. (2005). The Nucleation and Maintenance of Heterochromatin by a Histone Deacetylase in Fission Yeast. Mol. Cell 20, 173–185. 10.1016/j.molcel.2005.10.002.

39. Moazed, D. (2011). Mechanisms for the Inheritance of Chromatin States. Cell 146, 510–518. 10.1016/j.cell.2011.07.013.

40. Audergon, P.N.C.B., Catania, S., Kagansky, A., Tong, P., Shukla, M., Pidoux, A.L., and Allshire, R.C. (2015). Restricted epigenetic inheritance of H3K9 methylation. Science 348, 132–135. 10.1126/science.1260638.

41. Zhang, K., Mosch, K., Fischle, W., and Grewal, S.I.S. (2008). Roles of the Clr4 methyltransferase complex in nucleation, spreading and maintenance of heterochromatin. Nat. Struct. Mol. Biol. 15, 381–388. 10.1038/nsmb.1406.

42. Ragunathan, K., Jih, G., and Moazed, D. (2015). Epigenetic inheritance uncoupled from sequence-specific recruitment. Science 348, 1258699. 10.1126/science.1258699.

43. Nakayama, J., Klar, A.J.S., and Grewal, S.I.S. (2000). A Chromodomain Protein, Swi6, Performs Imprinting Functions in Fission Yeast during Mitosis and Meiosis. Cell 101, 307–317. 10.1016/s0092-8674(00)80840-5.

44. Volpe, T.A., Kidner, C., Hall, I.M., Teng, G., Grewal, S.I.S., and Martienssen, R.A. (2002). Regulation of Heterochromatic Silencing and Histone H3 Lysine-9 Methylation by RNAi. Science 297, 1833–1837. 10.1126/science.1074973.

45. Hall, I.M., Shankaranarayana, G.D., Noma, K. ichi, Ayoub, N., Cohen, A., and Grewal, S.I.S. (2002). Establishment and maintenance of a heterochromatin domain. Science 297, 2232– 2237. 10.1126/science.1076466.

46. Wang, X., and Moazed, D. (2017). DNA sequence-dependent epigenetic inheritance of gene silencing and histone H3K9 methylation. Science 356, 88–91. 10.1126/science.aaj2114.

47. Roguev, A., Schaft, D., Shevchenko, A., Aasland, R., Shevchenko, A., and Stewart, A.F. (2003). High Conservation of the Set1/Rad6 Axis of Histone 3 Lysine 4 Methylation in Budding and Fission Yeasts. J. Biol. Chem. 278, 8487–8493. 10.1074/jbc.M209562200.

48. Greenstein, R.A., Barrales, R.R., Sanchez, N.A., Bisanz, J.E., Braun, S., and Al-Sady, B. (2020). Set1/COMPASS repels heterochromatin invasion at euchromatic sites by disrupting Suv39/Clr4 activity and nucleosome stability. Genes Dev. 34, 99–117. 10.1101/gad.328468.119.

49. Flury, V., Georgescu, P.R., Iesmantavicius, V., Shimada, Y., Kuzdere, T., Braun, S., and Bühler, M. (2017). The Histone Acetyltransferase Mst2 Protects Active Chromatin from Epigenetic Silencing by Acetylating the Ubiquitin Ligase Brl1. Mol. Cell 67, 294–307.e9. 10.1016/j.molcel.2017.05.026.

50. Reddy, B.D., Wang, Y., Niu, L., Higuchi, E.C., Marguerat, S.B., Bähler, J., Smith, G.R., and Jia, S. (2011). Elimination of a specific histone H3K14 acetyltransferase complex bypasses the RNAi pathway to regulate pericentric heterochromatin functions. Genes Dev. 25, 214–219. 10.1101/gad.1993611.

51. Allshire, R.C., Javerzat, J.-P., Redhead, N.J., and Cranston, G. (1994). Position Effect Variegation at Fission Yeast Centromeres. Cell 76, 157–169. 10.1016/0092-8674(94)90180-5.

52. Zofall, M., Sandhu, R., Holla, S., Wheeler, D., and Grewal, S.I.S. (2022). Histone deacetylation primes self-propagation of heterochromatin domains to promote epigenetic inheritance. Nat. Struct. Mol. Biol. 29, 898–909. 10.1038/s41594-022-00830-7.

53. Shipkovenska, G., Durango, A., Kalocsay, M., Gygi, S.P., and Moazed, D. (2020). A conserved RNA degradation complex required for spreading and epigenetic inheritance of heterochromatin. eLife 9, e54341. 10.7554/eLife.54341.

54. Mikheyeva, I.V., Grady, P.J.R., Tamburini, F.B., Lorenz, D.R., and Cam, H.P. (2014). Multifaceted Genome Control by Set1 Dependent and Independent of H3K4 Methylation and the Set1C/COMPASS Complex. PLoS Genet. 10, e1004740. 10.1371/journal.pgen.1004740.

55. Svensson, J.P., Shukla, M., Menendez-Benito, V., Norman-Axelsson, U., Audergon, P., Sinha, I., Tanny, J.C., Allshire, R.C., and Ekwall, K. (2015). A nucleosome turnover map reveals that the stability of histone H4 Lys20 methylation depends on histone recycling in transcribed chromatin. Genome Res. 25, 872–883. 10.1101/gr.188870.114.

56. Wang, J., Reddy, B.D., and Jia, S. (2015). Rapid epigenetic adaptation to uncontrolled heterochromatin spreading. eLife 4, e06179. 10.7554/eLife.06179.

57. Sadaie, M., Kawaguchi, R., Ohtani, Y., Arisaka, F., Tanaka, K., Shirahige, K., and Nakayama, J.-i. (2008). Balance between Distinct HP1 Family Proteins Controls Heterochromatin Assembly in Fission Yeast. Mol. Cell. Biol. 28, 6973–6988. 10.1128/mcb.00791-08.

58. Creamer, K.M., Job, G., Shanker, S., Neale, G.A., Lin, Y.-C., Bartholomew, B., and Partridge, J.F. (2014). The Mi-2 Homolog Mit1 Actively Positions Nucleosomes within Heterochromatin To Suppress Transcription. Mol. Cell. Biol. 34, 2046–2061. 10.1128/MCB.01609-13.

59. Job, G., Brugger, C., Xu, T., Bañ Os Sanz, J.I., Partridge, J.F., and Schalch Correspondence, T. (2016). SHREC Silences Heterochromatin via Distinct Remodeling and Deacetylation Modules. Mol. Cell 62, 207–221. 10.1016/j.molcel.2016.03.016.

60. Allshire, R.C., Nimmo, E.R., Ekwall, K., Javerzat, J.P., and Cranston, G. (1995). Mutations derepressing silent centromeric domains in fission yeast disrupt chromosome segregation. Genes Dev. 9, 218–233. 10.1101/gad.9.2.218.

61. Grewal, S.I.S., and Klar, A.J.S. (1996). Chromosomal inheritance of epigenetic states in fission yeast during mitosis and meiosis. Cell 86, 95–101. 10.1016/S0092-8674(00)80080-X.

62. Roche, B., Arcangioli, B., and Martienssen, R.A. (2016). RNA interference is essential for cellular quiescence. Science 354, aah5651. 10.1126/science.aah5651.

63. Stirpe, A., Guidotti, N., Northall, S., Kilic, S., Hainard, A., Vadas, O., Fierz, B., and Schalch, T. (2020). SUV39 SET domains mediate crosstalk of heterochromatic histone marks Title: Regulation of SET domains by H3K14ub. BioRxiv Prepr. 10.1101/2020.06.30.177071.

64. Iglesias, N., Currie, M.A., Jih, G., Paulo, J.A., Siuti, N., Kalocsay, M., Gygi, S.P., and Moazed, D. (2018). Automethylation-induced conformational switch in Clr4 (Suv39h) maintains epigenetic stability. Nature 560, 504–508. 10.1038/s41586-018-0398-2.

65. Cutter DiPiazza, A.R., Taneja, N., Dhakshnamoorthy, J., Wheeler, D., Holla, S., and Grewal, S.I.S. (2021). Spreading and epigenetic inheritance of heterochromatin require a critical density of histone H3 lysine 9 tri-methylation. Proc. Natl. Acad. Sci. U. S. A. 118, 1–10. 10.1073/pnas.2100699118.

66. Trewick, S.C., Minc, E., Antonelli, R., Urano, T., and Allshire, R.C. (2007). The JmjC domain protein Epe1 prevents unregulated assembly and disassembly of heterochromatin. EMBO J. 26, 4670–4682. 10.1038/sj.emboj.7601892.

67. Holla, S., Dhakshnamoorthy, J., Folco, H.D., Wheeler, D., Zofall, M., Grewal Correspondence, S.I.S., Balachandran, V., Xiao, H., Sun, L.-L., and Grewal, S.I.S. (2020). Positioning Heterochromatin at the Nuclear Periphery Suppresses Histone Turnover to Promote Epigenetic Inheritance. Cell 180, 150–164. 10.1016/j.cell.2019.12.004.

68. Grewal, S.I.S. (2023). The molecular basis of heterochromatin assembly and epigenetic inheritance. Mol. Cell 83, 1767–1785. 10.1016/j.molcel.2023.04.020.

69. Stanton, B.Z., Chory, E.J., and Crabtree, G.R. (2018). Chemically induced proximity in biology and medicine. Science 359, eaao5902. 10.1126/science.aao5902.

70. Kagansky, A., Folco, H.D., Almeida, R., Pidoux, A.L., Boukaba, A., Simmer, F., Urano, T., Hamilton, G.L., and Allshire, R.C. (2009). Synthetic Heterochromatin Bypasses RNAi and Centromeric Repeats to Establish Functional Centromeres. Science 324, 1716–1719. 10.1126/science.1172026.

71. Erdel, F., Rademacher, A., Vlijm, R., Tünnermann, J., Frank, L., Weinmann, R., Schweigert, E., Yserentant, K., Hummert, J., Bauer, C., et al. (2020). Mouse Heterochromatin Adopts Digital Compaction States without Showing Hallmarks of HP1-Driven Liquid-Liquid Phase Separation. Mol. Cell 78, 1–14. 10.1016/j.molcel.2020.02.005.

72. Mizuguchi, T., Fudenberg, G., Mehta, S., Belton, J.M., Taneja, N., Folco, H.D., FitzGerald, P., Dekker, J., Mirny, L., Barrowman, J., et al. (2014). Cohesin-dependent globules and heterochromatin shape 3D genome architecture in S. pombe. Nature 516, 432–435. 10.1038/nature13833.

73. Owen, J.A., Osmanović, D., and Mirny, L.A. (2022). Design principles of 3D epigenetic memory systems. BioRxiv Prepr. 10.1101/2022.09.24.509332.

74. Abdulla, A.Z., Vaillant, C., and Jost, D. (2022). Painters in chromatin: a unified quantitative framework to systematically characterize epigenome regulation and memory. Nucleic Acids Res. 50, 9083–9104. 10.1093/nar/gkac702.

75. Bähler, J., Wu, J.-Q., Longtine, M.S., Shah, N.G., Mckenzie III, A., Steever, A.B., Wach, A., Philippsen, P., and Pringle, J.R. (1998). Heterologous modules for efficient and versatile PCR-based gene targeting inSchizosaccharomyces pombe. Yeast 14, 943–951. 10.1002/(SICI)1097-0061(199807)14:10<943::AID-YEA292>3.0.CO;2-Y.

76. Torres-Garcia, S., Di Pompeo, L., Eivers, L., Gaborieau, B., White, S.A., Pidoux, A.L., Kanigowska, P., Yaseen, I., Cai, Y., and Allshire, R.C. (2020). SpEDIT: A fast and efficient CRISPR/Cas9 method for fission yeast. Wellcome Open Res. 5, 274. 10.12688/wellcomeopenres.16405.1.

77. Raiymbek, G., An, S., Khurana, N., Gopinath, S., Larkin, A., Biswas, S., Trievel, R.C., Cho, U., and Ragunathan, K. (2020). An H3K9 methylation dependent protein interaction regulates the non-enzymatic function of a putative histone demethylase. eLife 9, e53155. 10.7554/elife.53155.

78. Li, H., and Durbin, R. (2010). Fast and accurate long-read alignment with Burrows–Wheeler transform. Bioinformatics 26, 589–595. 10.1093/bioinformatics/btp698.

79. Bolger, A.M., Lohse, M., and Usadel, B. (2014). Trimmomatic: a flexible trimmer for Illumina sequence data. Bioinformatics 30, 2114–2120. 10.1093/bioinformatics/btu170.

80. Zhang, Y., Liu, T., Meyer, C.A., Eeckhoute, J., Johnson, D.S., Bernstein, B.E., Nusbaum, C., Myers, R.M., Brown, M., Li, W., et al. (2008). Model-based Analysis of ChIP-Seq (MACS). Genome Biol. 9, R137. 10.1186/gb-2008-9-9-r137.

81. Ramírez, F., Ryan, D.P., Grüning, B., Bhardwaj, V., Kilpert, F., Richter, A.S., Heyne, S., Dündar, F., and Manke, T. (2016). deepTools2: a next generation web server for deep-sequencing data analysis. Nucleic Acids Res. 44, W160–W165. 10.1093/nar/gkw257.

82. Dobin, A., Davis, C.A., Schlesinger, F., Drenkow, J., Zaleski, C., Jha, S., Batut, P., Chaisson, M., and Gingeras, T.R. (2013). STAR: ultrafast universal RNA-seq aligner. Bioinformatics 29, 15–21. 10.1093/bioinformatics/bts635.

83. Love, M.I., Huber, W., and Anders, S. (2014). Moderated estimation of fold change and dispersion for RNA-seq data with DESeq2. Genome Biol. 15, 550. 10.1186/s13059-014-0550-8.

