## Supplemental Figures and Tables for "Uncoupling the distinct functions of HP1 proteins during heterochromatin establishment and maintenance"

**Figure S1. Related to Figure 1.**

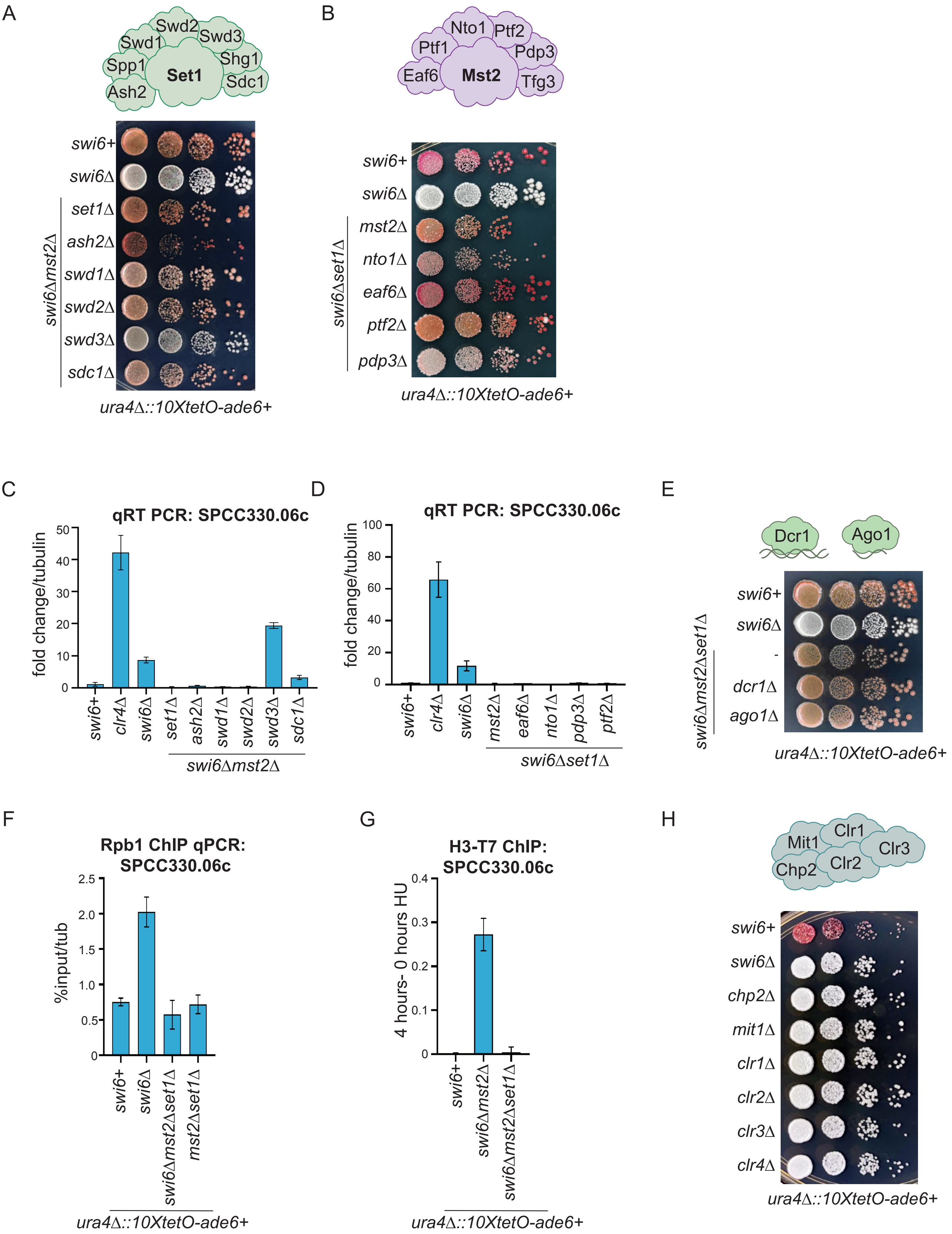

**Figure S1. Swi6 can be bypassed in transcriptional gene silencing by deleting subunits of the Set1 complex or the Mst2 complex.** (**A**) Silencing assay of *ura4Δ::10XtetO-ade6+* reporter strains with deletions of different Set1 complex subunits. (**B**) Silencing assay of *ura4Δ::10XtetO-ade6+* reporter strains with deletions of different Mst2 complex subunits. (**C**,**D**) qRT-PCR measurements of RNA levels of *SPCC330.06c* in each indicated genotype. Error bars indicate SD (N=2) (**E**) Silencing assay measuring Swi6 independent silencing in RNAi deletion strains. (**F**) ChIP-qPCR to measure the main subunit of RNA polymerase II, Rpb1, in each indicated genotype (N=2). (**G**) ChIP-qPCR of newly incorporated histone H3-T7 at *SPCC330.06c* in the indicated genotypes. Input normalized ChIP values of uninduced samples (0h) were subtracted from input normalized ChIP values of β-estradiol induced samples (4h). (**H**) Silencing assay when in the absence of subunits of SHREC.

### Figure S2.

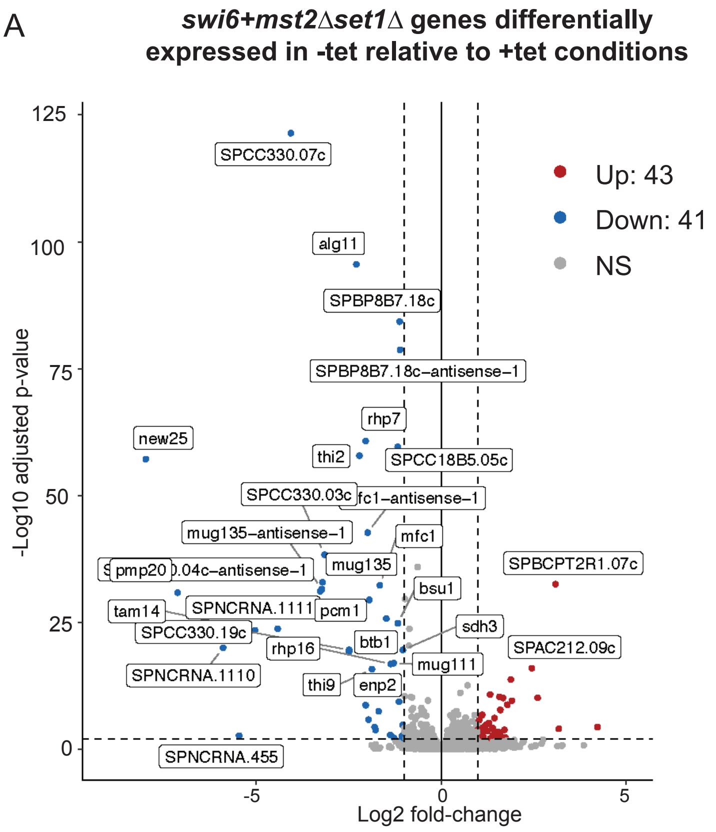
A
swi6+mst2Δset1Δ genes differentially expressed in -tet relative to +tet conditions
Up: 43
Down: 41
NS
-Log10 adjusted p-value
Log2 fold-change
125
100
75
50
25
0
-5
0
5
SPCC330.07c
alg11
SPBP8B7.18c
SPBP8B7.18c–antisense–1
rhp7
new25
thi2
SPCC18B5.05c
SPCC330.03c
mug135–antisense–1
mfc1
mug135
pmp20
bsu1
SPBCPT2R1.07c
tam14
SPNCRNA.1111
pcm1
sdh3
SPCC330.19c
rhp16
btb1
SPAC212.09c
mug111
SPNCRNA.1110
thi9
enp2
SPNCRNA.455

**Figure S2. Downregulation of gene expression relies on Clr4 tethering when Set1 and Mst2 are deleted.** (**A**) Volcano plots of RNA levels in -tet media relative to +tet media in *swi6+mst2∆set1∆* cells. Genes proximal to the *10xtetO-ade6+* reporter are labelled in the volcano plot.

**Figure S3. Related to Figure 3.**

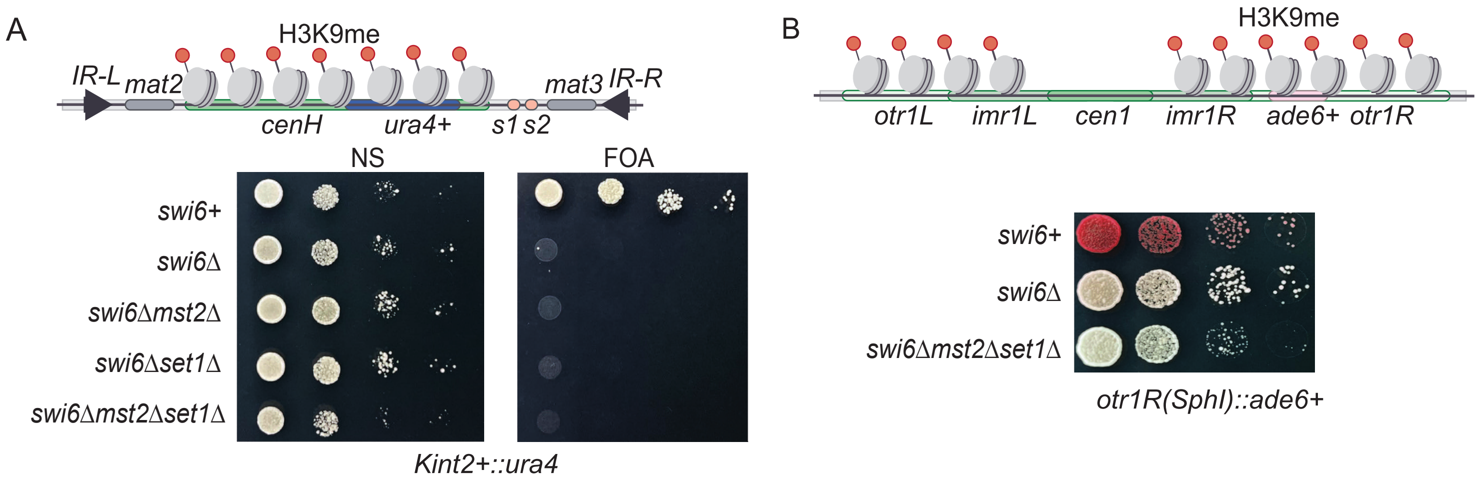

**Figure S3. Swi6 is not bypassed at the endogenous *mat* locus or pericentromeric repeats when Set1 and Mst2 are deleted.** (**A**) Schematic of the *mat* locus with a ura4+ reporter and intact RNAi dependent *cenH* sequence. Silencing assay of *ura4+* reporter in the indicated genotypes. (**B**) Silencing of the pericentromeric *otr1R(SphI):ade6+* reporter in the indicated genotypes.

**Figure S4. Related to Figure 6.**

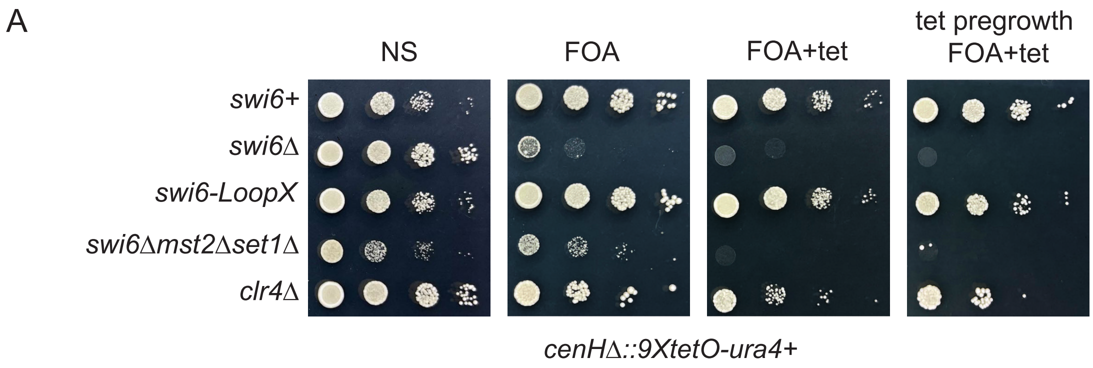

**Figure S4. Swi6 oligomerization is not required for maintenance at the *mat* reporter locus.** (**A**) Silencing assay of *cenHΔ::9XtetO-ura4+* reporter strains in the indicated genotypes.

**Table S1.**
**Strains used in this study**

| | | |
|---|---|---|
| KR18 | *h-, leu1-32, ade6-M210, ura4Δ::10XTetO-ade6, clr4+, trp1:nat-clr4p-TetR-clr4ΔCD* | Moazed lab |
| KR53 | *h-, leu1-32, ade6-M210, ura4Δ::10XTetO-ade6, clr4+, trp1:nat-clr4p-TetR-clr4ΔCD, swi6Δ::hphMX6* | Moazed lab |
| KR1303 | *h-, leu1-32, ade6-M210, ura4Δ::10XTetO-ade6, clr4+, trp1:nat-clr4p-TetR-clr4ΔCD, swi6Δ::hphMX6, set1Δ::bsd, mst2Δ::ura4* | This study |
| KR1135 | *h-, leu1-32, ade6-M210, ura4Δ::10XTetO-ade6, clr4+, trp1:nat-clr4p-TetR-clr4ΔCD, swi6Δ::hphMX6, set1Δ::kanMX6* | This study |
| KR1433 | *h-, leu1-32, ade6-M210, ura4Δ::10XTetO-ade6, clr4+, trp1:nat-clr4p-TetR-clr4ΔCD, swi6Δ::hphMX6 mst2Δ::ura4* | This study |
| KR2015 | *h-, leu1-32, ade6-M210, ura4Δ::10XTetO-ade6, clr4+, trp1:nat-clr4p-TetR-clr4ΔCD, set1Δ:bsd mst2Δ::ura4* | This study |
| KR24 | *h-, leu1-32, ade6-M210, ura4Δ::10XTetO-ade6, clr4Δ::kanMX6* | Moazed lab |
| KR38 | *h- leu1-32 ade6-M210 ura4Δ::10XTetO-ura4p(114bp)-ura4-GFP* | Moazed lab |
| KR2043 | *h- leu1-32 ade6-M210 ura4Δ::10XTetO-ura4p(114bp)-ura4-GFP, swi6Δ::hphMX6* | This study |
| KR2049 | *h- leu1-32 ade6-M210 ura4Δ::10XTetO-ura4p(114bp)-ura4-GFP, swi6Δ::hphMX6, mst2Δ::kanMX6* | This study |
| KR2051 | *h- leu1-32 ade6-M210 ura4Δ::10XTetO-ura4p(114bp)-ura4-GFP, swi6Δ::hphMX6, set1Δ::bsd* | This study |
| KR2056 | *h- leu1-32 ade6-M210 ura4Δ::10XTetO-ura4p(114bp)-ura4-GFP, swi6Δ::hphMX6, set1Δ::bsd, mst2Δ::kanMX6* | This study |
| KR1714 | *h-, leu1-32, ade6-M210, ura4Δ::10XTetO-ade6, clr4+, trp1:nat-clr4p-TetR-clr4ΔCD, swi6Δ::kanMX6, set1Δ::bsd, mst2Δ::ura4, chp2Δ::hphMX6* | This study |
| KR1768 | *h-, leu1-32, ade6-M210, ura4Δ::10XTetO-ade6, clr4+, trp1:nat-clr4p-TetR-clr4ΔCD, swi6Δ::hphMX6, set1Δ::bsd, mst2Δ::ura4, mit1Δ::kanMX6* | This study |
| KR2150 | *h-, leu1-32, ade6-M210, ura4Δ::10XTetO-ade6, clr4+, trp1:nat-clr4p-TetR-clr4ΔCD, swi6Δ::hphMX6, set1Δ::bsd, mst2Δ::ura4, clr1Δ::kanMX6* | This study |
| KR2152 | *h-, leu1-32, ade6-M210, ura4Δ::10XTetO-ade6, clr4+, trp1:nat-clr4p-TetR-clr4ΔCD, swi6Δ::hphMX6, set1Δ::bsd, mst2Δ::ura4, clr2Δ::kanMX6* | This study |
| KR54 | *h-, leu1-32, ade6-M210, ura4Δ::10XTetO-ade6, clr4+, trp1:nat-clr4p-TetR-clr4ΔCD, chp2Δ::hphMX6* | Moazed lab |
| KR1744 | *h+, leu1-32, ade6-M210, ura4Δ::10XTetO-ade6, clr4+, trp1:nat-clr4p-TetR-clr4ΔCD, chp2Δ::hphMX6, mst2Δ::ura4, set1Δ::bsd* | This study |
| KR1435 | *h-, leu1-32, ade6-M210, ura4Δ::10XTetO-ade6, clr4+, trp1:nat-clr4p-TetR-clr4ΔCD, chp2Δ::hphMX6, mst2Δ::ura4* | This study |
| KR1406 | *h+, leu1-32, ade6-M210, ura4Δ::10XTetO-ade6, clr4+, trp1:nat-clr4p-TetR-clr4ΔCD, chp2Δ::hphMX6, set1Δ::bsd* | This study |
| KR2175 | *h-, leu1-32, ade6-M210, ura4Δ::10XTetO-ade6, clr4+, trp1:nat-clr4p-TetR-clr4ΔCD, swi6Δ::hphMX6 mst2Δ::ura4, ash2Δ::kanMX6* | This study |
| KR2033 | *h-, leu1-32, ade6-M210, ura4Δ::10XTetO-ade6, clr4+, trp1:nat-clr4p-TetR-clr4ΔCD, swi6Δ::hphMX6 mst2Δ::ura4, swd1Δ::kanMX6* | This study |

| | | |
|---|---|---|
| KR2035 | *h-, leu1-32, ade6-M210, ura4∆::10XTetO-ade6, clr4+, trp1:nat-clr4p-TetR-clr4∆CD, swi6∆::hphMX6 mst2∆::ura4, swd2∆::kanMX6* | This study |
| KR2094 | *h-, leu1-32, ade6-M210, ura4∆::10XTetO-ade6, clr4+, trp1:nat-clr4p-TetR-clr4∆CD, swi6∆::hphMX6 mst2∆::ura4, swd3∆::bsd* | This study |
| KR2096 | *h-, leu1-32, ade6-M210, ura4∆::10XTetO-ade6, clr4+, trp1:nat-clr4p-TetR-clr4∆CD, swi6∆::hphMX6 mst2∆::ura4, sdc1∆::bsd* | This study |
| KR2036 | *h-, leu1-32, ade6-M210, ura4∆::10XTetO-ade6, clr4+, trp1:nat-clr4p-TetR-clr4∆CD, swi6∆::hphMX6, set1∆::kanMX6, nto1∆::bsd* | This study |
| KR2097 | *h-, leu1-32, ade6-M210, ura4∆::10XTetO-ade6, clr4+, trp1:nat-clr4p-TetR-clr4∆CD, swi6∆::hphMX6, set1∆::kanMX6, eaf6∆::bsd* | This study |
| KR2032 | *h-, leu1-32, ade6-M210, ura4∆::10XTetO-ade6, clr4+, trp1:nat-clr4p-TetR-clr4∆CD, swi6∆::hphMX6, set1∆::kanMX6, ptf2∆::bsd* | This study |
| KR2031 | *h-, leu1-32, ade6-M210, ura4∆::10XTetO-ade6, clr4+, trp1:nat-clr4p-TetR-clr4∆CD, swi6∆::hphMX6, set1∆::kanMX6, pdp3∆::bsd* | This study |
| KR2208 | *h+, leu1-32, ade6-M210, ura4∆::10XTetO-ade6, clr4+, trp1:nat-clr4p-TetR-clr4∆CD, swi6∆::hphMX6, set1∆::bsd, mst2∆::ura4, dcr1∆::kanMX6* | This study |
| KR2212 | *h+, leu1-32, ade6-M210, ura4∆::10XTetO-ade6, clr4+, trp1:nat-clr4p-TetR-clr4∆CD, swi6∆::hphMX6, set1∆::bsd, mst2∆::ura4, ago1∆::kanMX6* | This study |
| KR578 | *h90 ade6+ leu1-32 ura4-D18, CenH∆::9xtetO-ura4, trp1:nat-clr4p-TetR-clr4ΔCD* | Moazed lab |
| KR1715 | *h90 ade6+ leu1-32 ura4-D18, CenH∆::9xtetO-ura4, trp1:nat-clr4p-TetR-clr4ΔCD, swi6∆::bsd* | This study |
| KR1819 | *h90 ade6+ leu1-32 ura4-D18, CenH∆::9xtetO-ura4, trp1:nat-clr4p-TetR-clr4ΔCD, swi6∆::hgh, mst2∆::kanMX6, set1∆::bsd* | This study |
| KR1757 | *h90 ade6+ leu1-32 ura4-D18, CenH∆::9xtetO-ura4, trp1:nat-clr4p-TetR-clr4ΔCD, swi6∆::hgh, mst2∆::kanMX6* | This study |
| KR1763 | *h90 ade6+ leu1-32 ura4-D18, CenH∆::9xtetO-ura4, trp1:nat-clr4p-TetR-clr4ΔCD, swi6∆::bsd, set1∆::kanMX6* | This study |
| KR683 | *h90 leu1-32 his2- ura4 DS/E ade6-M210 Kint2::ura4+* | Jia lab |
| KR1719 | *h90 leu1-32 his2- ura4 DS/E ade6-M210 Kint2::ura4+, swi6∆::bsd* | This study |
| KR1762 | *h90 leu1-32 his2- ura4 DS/E ade6-M210 Kint2::ura4+, swi6∆::bsd, mst2∆::hgh* | This study |
| KR2374 | *h90 leu1-32 his2- ura4 DS/E ade6-M210 Kint2::ura4+, swi6∆::bsd, set1∆::kanMX6* | This study |
| KR2371 | *h90 leu1-32 his2- ura4 DS/E ade6-M210 Kint2::ura4+, swi6∆::bsd, set1∆::kanMX6, mst2∆::hph* | This study |
| KR343 | *h+, otr1R(SphI)::ade6+, ura4-D18, leu1-32, ade6-M210* | Moazed lab |
| KR450 | *h+, otr1R(SphI)::ade6+, ura4-D18, leu1-32, ade6-M210, swi6∆::nat* | Moazed lab |
| KR1723 | *h-, otr1R(SphI)::ade6+, ura4-D18, leu1-32, ade6-M210, swi6∆::nat, mst2∆::ura4, set1∆::kanMX6* | This study |
| KR33 | *h-, leu1-32, ade6?, ura4∆::10XTetO-ade6, clr4+, trp1:nat-clr4p-TetR-clr4∆CD, epe1∆::kanMX6* | Moazed lab |
| KR59 | *h-, leu1-32, ade6?, ura4∆::10XTetO-ade6, clr4+, trp1:nat-clr4p-TetR-clr4∆CD, epe1∆::kanMX6, swi6∆::hph* | Moazed lab |
| KR1753 | *h- leu1-32 ade6? ura4∆::10XTetO-ade6 clr4+ trp1:nat-clr4p-TetR-clr4∆CD, epe1∆::kanMX6 swi6∆::hphMX6 set1∆:bsd mst2∆::ura4* | This study |

| | | |
|---|---|---|
| KR2643 | *h-, SPY5071 clr4+, trp1:nat-clr4p-TetR-clr4ΔCD, hht1-G13D* | This study |
| KR2646 | *h-, SPY5071 clr4+, trp1:nat-clr4p-TetR-clr4ΔCD, hht1-G13D, swi6Δ::hph* | This study |
| KR2693 | *h-, SPY5071 clr4+, trp1:nat-clr4p-TetR-clr4ΔCD, hht1-G13D, swi6Δ::hph, set1Δ::bsd, mst2Δ::ura4 #1* | This study |
| KR2694 | *h-, SPY5071 clr4+, trp1:nat-clr4p-TetR-clr4ΔCD, hht1-G13D, swi6Δ::hph, set1Δ::bsd, mst2Δ::ura4 #2* | This study |
| KR2762 | *h-, SPY5071 clr4+, trp1:nat-clr4p-TetR-clr4ΔCD, hht1-G13D, set1Δ::bsd, mst2Δ::ura4* | This study |
| KR2781 | *h- leu1-32 ade6? ura4Δ::10XTetO-ade6 clr4+, trp1:nat-clr4p-TetR-clr4DCD , epe1Δ::kanMX6, swi6-ARK* | This study |
| KR2783 | *h90 ade6+ leu1-32 ura4-D18, CenHΔ::9xtetO-ura4, trp1:nat-clr4p-TetR-clr4ΔCD, swi6-ARK* | This study |
| KR2973 | *h+, ars1::prad15 cre-EBD-LEU2, H3.2 lox-HA-HYG-lox-T7, cdc25+, leu1-32, ura4Δ::10XTetO-ade6, clr4+, trp1:nat-clr4p-TetR-clr4ΔCD* | This study |
| KR2974 | *h-, ars1::prad15 cre-EBD-LEU2, H3.2 lox-HA-HYG-lox-T7, cdc25+, leu1-32, ura4Δ::10XTetO-ade6, clr4+, trp1:nat-clr4p-TetR-clr4ΔCD, swi6Δ::kanMX* | This study |
| KR2975 | *h-, ars1::prad15 cre-EBD-LEU2, H3.2 lox-HA-HYG-lox-T7, cdc25+, leu1-32, ura4Δ::10XTetO-ade6, clr4+, trp1:nat-clr4p-TetR-clr4ΔCD, swi6Δ::kanMX mst2Δ::ura4* | This study |

**Table S2.**
**Primers used in this study**

| | |
|---|---|
| ura4 F | CGTGGTCTCTTGCTTTTGGC |
| ura4 R | CATCCAAGCCGATACCAGGG |
| ade6 F | CAAACCCTTGCCATGGATGC |
| ade6 R | ACCTGAATTGTGAGGCCGAG |
| SPCC330.06c F | GCCGTAAATGACGTTTTCGTCACC |
| SPCC330.06c R | ACCTTGACAACCTTGCCATTCTCG |
| GFP F | ACCCAGACCACATGAAGCAACAC |
| GFP R | CTTGTGACCCAAGATGTTACCGTC |
| tub F | AACGCTTGGCCATGGAATACACG |
| tub R | GAGAGGCGGTGATGGAAGAAACAAC |
